# Corticotropin releasing factor alters the functional diversity of accumbal cholinergic interneurons

**DOI:** 10.1101/2023.09.17.558116

**Authors:** Anna E. Ingebretson, Yanaira Alonso-Caraballo, John A. Razidlo, Julia C. Lemos

## Abstract

Cholinergic interneurons (ChIs) provide the main source of acetylcholine in the striatum and have emerged as a critical modulator of behavioral flexibility, motivation, and associative learning. In the dorsal striatum, ChIs display heterogeneous firing patterns. Here, we investigated the spontaneous firing patterns of ChIs in the nucleus accumbens (NAc) shell, a region of the ventral striatum. We identified four distinct ChI firing signatures: regular single-spiking, irregular single-spiking, rhythmic bursting, and a mixed-mode pattern composed of bursting activity and regular single spiking. ChIs from females had lower firing rates compared to males and had both a higher proportion of mixed-mode firing patterns and a lower proportion of regular single-spiking neurons compared to males. We further observed that across the estrous cycle, the diestrus phase was characterized by higher proportions of irregular ChI firing patterns compared to other phases. Using pooled data from males and females, we examined how the stress-associated neuropeptide corticotropin releasing factor (CRF) impacts these firing patterns. ChI firing patterns showed differential sensitivity to CRF. This translated into differential ChI sensitivity to CRF across the estrous cycle. Furthermore, CRF shifted the proportion of ChI firing patterns toward more regular spiking activity over bursting patterns. Finally, we found that repeated stressor exposure altered ChI firing patterns and sensitivity to CRF in the NAc core, but not the NAc shell. These findings highlight the heterogeneous nature of ChI firing patterns, which may have implications for accumbal-dependent motivated behaviors.

**New and Noteworthy:** ChIs within the dorsal and ventral striatum can exert a huge influence on network output and motivated behaviors. However, the firing patterns and neuromodulation of ChIs within the ventral striatum, specifically the NAc shell, are understudied. Here we report that NAc shell ChIs have heterogenous ChI firing patterns that are labile and can be modulated by the stress-linked neuropeptide CRF and by the estrous cycle.

## Introduction

Cholinergic interneurons (ChIs) provide the main source of acetylcholine modulation of striatal circuit function (Gonzales and Smith, 2015). ChIs make up approximately 1-2% of all the neurons within the dorsal and ventral striatum (Bolam, Wainer and Smith, 1984; Bonsi et al., 2011; Nosaka and Wickens, 2022). However, these large aspiny neurons are highly ramified such that this relatively small number of neurons can serve as a master regulator of striatal output (Gonzales and Smith, 2015; Nosaka and Wickens, 2022). A central functional property of this population of neurons is that they are spontaneously active both in in vivo and ex vivo preparations (Gonzales and Smith, 2015). In the dorsal striatum, ChIs display heterogeneous firing patterns. Bennett and Wilson (1999) identified three categories of spontaneously firing ChIs within the dorsal striatum. “Regular,” “irregular,” and “rhythmic bursting” signatures could be distinguished by their differential firing rates and coefficients of variation of the inter-event interval (IEI) distribution (Bennett and Wilson, 1999). “Regular” ChIs had high firing rates distinguished by regularly occurring single spikes that had narrow, unimodal IEI distributions and low IEI CVs, characteristic of canonical pacemaker firing. “Irregular” ChIs had lower firing rates and higher CVs, indicating greater variability in spike timing. Finally, “rhythmic bursting” ChIs were characterized by a rhythmic firing pattern consisting of bursts of spikes interspersed with pauses in firing activity that yielded a skewed or bimodal IEI distribution with shorter intraburst intervals comprising the mean interval and longer interburst pauses forming the tail of the distribution; this high degree of temporal variability produced low overall firing rates and very high IEI CVs (Bennett, Callaway and Wilson, 2000; Bennett and Wilson, 1999). Bennet and Wilson went on to demonstrate that these disparate firing properties were not driven by excitatory or inhibitory synaptic transmission, but by differential intrinsic properties. Specifically, the small calcium-activated potassium channel (sK) was identified as a key regulator of firing properties (Bennett, Callaway and Wilson, 2000; Bennett and Wilson, 1999; Goldberg and Wilson, 2005; Wilson, 2005).

In contrast to the dorsal striatum, the firing properties of different cell types within the NAc core and shell, including ChIs, remain largely unexplored. It is unknown whether the functional heterogeneity of ChIs in the dorsal striatum is also present in the ventral striatum. While early studies included both male and female animals, sex and sex hormones as biological variables of interest were not explored. Furthermore, while it has been shown that manipulations that reduce or disrupt normal ChI firing cause depression-like behavior in rodents, the impact of stressor exposure on ChI firing patterns or properties has not been explored (Cheng et al., 2019; Hanada et al., 2018; Warner-Schmidt et al., 2012). In addition, the functional relevance of the complex spiking behavior of ChIs has not been fully examined. Understanding the cellular properties of NAc shell ChIs may provide important mechanistic insights into their role in motivated behavior.

Recent work has renewed interest in the cellular properties of ChIs and how heterogeneous firing modes might relate to the functional role of ChIs in motivated behavior. For example, shifts from tonic ChI pacemaker activity to a burst-pause mode in the dorsal striatum has been suggested to encode a salient environmental stimulus that generates an attentional and, perhaps, a motivational shift (Ding et al., 2010; Krok et al., 2023; Nosaka and Wickens, 2022; Zhang and Cragg, 2017; Zucca et al., 2018). To our knowledge, no in vivo electrophysiology studies have been carried out focusing on ChIs in the NAc. However, both bursts and ramps of ChI population activity are correlated with motivated approach behavior in the NAc (Mohebi, Collins and Berke, 2023). Furthermore, ChI population activity within the NAc shell both tracks with and promotes reward learning (Al-Hasani et al., 2021). While a mechanistic understanding of how these salient environmental stimuli are translated into molecular and cellular signals in the NAc is still unclear, one potential mechanism is through the release of neuropeptides.

Corticotropin-releasing factor (CRF) is a stress-associated neuropeptide that is widely distributed and released in the brain and the periphery during periods of high arousal and salience, including novelty (Bale and Vale, 2004; Dabrowska et al., 2013; Henckens, Deussing and Chen, 2016; Lemos et al., 2012). In the cortex, CRF is released in response to anticipatory cues that signal food delivery (Merali, McIntosh and Anisman, 2004). In the NAc, CRF is released in response to novelty and facilitates novelty exploration (Lemos et al., 2012). Exogenous application of CRF in the NAc can promote appetitive behaviors and potentiate both dopamine and acetylcholine transmission (Chen et al., 2012; Lemos et al., 2012; Lim et al., 2007; Pecina, Schulkin and Berridge, 2006). Finally, CRF potentiates ChI firing rate in the NAc core and dorsal striatum via activation of CRF type 1 receptors (CRF-R1), cAMP, and sK channel activation in male mice (Lemos, Shin and Alvarez, 2019). Given the role for striatal ChIs in encoding salient stimuli, CRF might shape salience encoding in the NAc via modulation of ChI firing patterns. In this study, we investigated the diversity of spontaneous firing properties in the NAc shell of both male and female mice and across the estrous cycle in female mice. We had previously not characterized ChI firing patterns or CRF effects in the NAc shell. We examined if ChIs with different spontaneous firing properties differentially respond to CRF application at different concentrations. During this investigation, we found that there were four distinct categories of ChI based on temporal firing patterns. We also found interesting sex differences and sensitivity to CRF based on this categorization. Importantly, we found that CRF shifts the distribution of firing modes toward a more regular tonic firing pattern as opposed to a more rhythmic or bursting mode. Finally, we examined how prior stress history impacts ChI firing patterns in the NAc core and shell and how it may shift CRF sensitivity. These data may provide a cellular mechanism for how salient stimuli shift the circuit level output of the NAc to facilitate appropriate behavioral responses to the environment.

## Methods and Material

All procedures were performed in accordance with guidelines from the Institutional Animal Care and Use Committee at the University of Minnesota.

### Animals

Male and female mice (post-natal day 60-180) were group housed and kept under a 12h-light cycle (6:30 ON/18:30 OFF) with food and water available *ad libitum*. For all ex vivo electrophysiology experiments, ChAT-IRES-Cre^+/-^ mice (B6N.129S6(B6)- Chattm2(cre)Lowl/J, Jackson Laboratory stock number 018957) crossed with Ai14;tdTomato reporter mice (B6.Cg-Gt(ROSA)26Sortm14(CAG-tdTomato)Hze/J, Jackson Laboratory stock number 007914) were used to easily visualize ChIs. C57BL6/J males were use for in situ hybridization experiments.

### Estrous cycle tracking

Vaginal lavage methods were used to track the estrous cycle of female mice over an 8–10 day period in order to capture two full cycles. We used two different methodologies that yielded the same conclusions. First, 20 µl of sterile saline was rapidly pipetted in and out of the vagina to collect vaginal cells within the solution. The sample was dried, stained with cresyl violet, and assessed under the microscope. Based on the proportion of leukocytes, cornified epithelial cells and nucleated epithelial cells, as well as the size of the vaginal opening, we classified females as in proestrus, estrus, metestrus or diestrus (Cora, Kooistra and Travlos, 2015). Later, we were able to use a 24 well plate and inverted light microscope without cresyl violet staining as this allowed for higher throughput (Alonso-Caraballo and Ferrario, 2019).

### Ex vivo electrophysiology

Coronal brain slices (240 µm) containing the NAc core and shell were prepared from 8-24-week-old *ChAT-ires-CRE^+/-^;Ai14 tdTomato* mice. Slices were cut in ice-cold cutting solution (in mM): 225 sucrose, 13.9 NaCl, 26.2 NaHCO_3_, 1 NaH_2_PO_4_, 1.25 glucose, 2.5 KCl, 0.1 CaCl_2_, 4.9 MgCl_2_ and 3 kynurenic acid. Slices were maintained in oxygenated artificial cerebrospinal fluid (ACSF) containing (in mM): 124 NaCl, 2.5 KCl, 2.5 CaCl_2_, 1.3 MgCl_2_, 26.2 NaHCO_3_, 1 NaH_2_PO_4_ and 20 glucose (∼310–315 mOsm) at room temperature following a 1hr recovery period at 33⁰C. Cell-attached recordings were made in voltage clamp mode at 30-33⁰C to assess firing properties and to measure the effect of CRF on ChI firing frequency without causing “run-down” due to dialyzing the cell with internal solution in a whole-cell configuration. For cell-attached recordings, electrodes were filled with filtered ACSF identical to the external solution. A gigaohm seal was achieved, maintained, and monitored. Recordings were conducted for a maximum of 15 minutes. Cells in which the gigaohm seal had degraded were excluded; seal degradation was determined by rapid changes in holding current past 100 pA. Data were acquired at 5 kHz and filtered at 1 kHz using either a SutterPatch or Multiclamp 700B (Molecular Devices). Data was analyzed using Igor or pClamp (Clampfit, v. 10.3).

### Drug application

2,3-Dioxo-6-nitro-1,2,3,4-tetrahydrobenzo[*f*]quinoxaline-7- sulfonamide (NBQX), 3-((*R*)-2-Carboxypiperazin-4-yl)-propyl-1-phosphonic acid (R- CPP), and rat/human CRF were acquired from R & D systems (Tocris) and bath applied to the slice preparation. NBQX and CPP were both applied at a final concentration of 5 µM. We chose to use ion channel blockers at submaximal concentrations since, in our experience, maximal concentrations of these blockers can lead to total shut down of firing and disruption of the gigaohm seal: ZD 7288 (HCN channel blocker, 500 nM); apamin (sK channel blocker, 150 nM), AP-4 (Kv1 channel blocker, 500 µM), Phrixotoxin- 1 (Kv4 blocker, 50 nM). CRF was bath applied at concentrations of 0, 3, 10, and 100 nM. These concentrations of CRF were based on our previous work (Lemos et al., 2019), where we applied CRF to NAc core slices from males and found that the EC50 was 8.6 nM and the maximal concentration was 100 nM (i.e. 300 nM did not produce a larger effect). The last three minutes of CRF application were averaged to produce a mean response.

### Fluorescent *in situ* hybridization (ISH) using RNAScope

Brains were rapidly dissected from male mice and flash frozen in isopentane on dry ice. Brains were kept in a -80°C freezer until they were sectioned. Coronal or sagittal brain slices (16 µm) containing the DS and NAc were thaw mounted onto Superfrost plus slides (Electron Microscopy Sciences) utilizing a Leica CM 1900 cryostat maintained at -20°C. Note, prior to section, brains were equilibrated in the cryostat for at least 2 hrs. Slides were cleaned with RNaseZap, a decontamination solution used to prevent mRNA degradation (Invitrogen). Slides were stored at -80°C.

RNAScope ISH was conducted according to the Advanced Cell Diagnostics user manual and as previously reported (Lemos, Shin and Alvarez, 2019). Briefly, slides were fixed in 10% neutral buffered formalin for 20 min at 4°C. Slides were washed 2x1 min with 1x PBS, before dehydration with 50% ethanol (1 x 5 min), 70% ethanol (1 x 5 min), and 100% ethanol (2 x 5 min). Slides were incubated in 100% ethanol at -20°C overnight. The following day, slides were dried at room temperature (RT) for 10 min. A hydrophobic barrier was drawn around the sections using a hydrophobic pen and allowed to dry for 15 min at RT. Sections were then incubated with Protease Pretreat-4 solution (Advanced Cell Diagnostics) for 20 min at RT. Slides were washed with ddH2O (2 x 1 min), before being incubated with the appropriate probes for 2 hr at 40°C in the HybEZ oven (Advanced Cell Diagnostics). The following probes were purchased from Advanced Cell Diagnostics: Mm-*Crh1*-C1 (ACD Cat no: 418011), Mm-*Crh*-C1 (ACD Cat no: 316091), and Mm-*Chat*-C2 (ACD Cat no:408731-C2). Following incubation with the appropriate probes, slides were subjected to a series of amplification steps at 40°C in the HybEZ oven with 2 x 2 min washes (with agitation) in between each amplification step at RT. Amplification steps were performed as follows: Amp 1 at 40°C for 30 min. Amp 2 at 40°C for 15 min. Amp 3 at 40°C for 30 min. Amp 4-Alt A at 40°C for 15 min. A DAPI-containing solution was applied to sections (one slide at a time) at RT for 20 sec. Finally, slides were coverslipped using ProLong Gold Antifade mounting media (Invitrogen) and stored at 4°C until imaging on a confocal microscope (Zeiss).

### Image analysis and quantification

Sections were imaged using a Zeiss confocal microscope and Zen software. Unique 20x (5 µm thick) and 40x images (2 µm thick) were acquired from the NAc of stress-naïve and stress-exposed mice using the same software and hardware settings. The settings were titrated for each specific experimental probe. Quantification was done using Fiji/ImageJ software. Numbers of DAPI cells were automatically generated using the particle counter function in ImageJ. Numbers of ChAT cells and ChAT+/CRF-R1 or +CRF positive cells were manually counted using the cell counter function. For both ChAT and DAPI cells, cells were considered positive for the experimental probe if there were >5 particles clustered around (but not in) the cell nucleus. Numbers of puncta per ChAT cell (67-71 unique cells for naïve and stress-exposed mice) were analyzed in a semi-automated fashion. Masks were generated using the ChAT as a reference point. Rather than using the outline of the cell itself, we used a uniform circle that was the approximate size of the soma and was kept consistent throughout the analysis. The fluorescent image of the experiment probe, in this case Mm- Crh1 (ChAT+) or Mm-Crh (DAPI), was converted into a binary image after being thresholded. The thresholding was kept consistent across images. Finally, the mask generated with the cell maker image was combined with the binary image of the experimental probe. This generated an image of only the puncta around the soma of our cells of interest. We then used the particle analyzer function to automatically count the number of puncta.

### Forced swim stress

Male mice were moved to a behavioral suite and allowed to acclimate for at least 30 min. The swim stress procedure was done as described in (Lemos et al., 2012). Briefly, animals were placed in a 20 cm cylinder filled with 4L of water maintained at 30 ± 1°C for 15 mins on Day 1 and 4 x 6 min on Day 2, separated by 6 min intervals in their home cages. Animals were returned to their home cages for ∼7 days (6-8 days) or ∼14 days (14-18 days), and then ex vivo slices are prepared. This swim stress procedure has been shown to produce CRF release and lead to long-term behavioral changes (Land et al., 2008). We used littermates that were maintained in their homecages and not exposed to stress or stress-exposed animals as our control group.

### Locomotor behavior

Animals were placed in a large (50 × 50 × 40 cm), novel circular open field for 60 minutes and video monitored. Noldus Ethovision software (v. 14) was used to analyze locomotor data.

### Statistics and classification

We ran a power analysis to determine sample sizes using G*Power 3.1 based on effect sizes identified previously (Lemos, Shin and Alvarez, 2019). The power analysis is based on the expected magnitudes and standard deviations of this previous study as well as an α-value of 0.05. Notably, our previous study using stress-naïve males had statistically less variability than females or stress-exposed animals and we had to increase our replicates as a result. Statistical analysis was performed in Prism (GraphPad) and Excel on spiking data extracted from the SutterPatch or Multiclamp 700B platforms. For each cell, analysis was performed on three minutes of stable, continuous recording following a three-minute equilibration period. The same was done following wash on and equilibration of CRF. Cells with firing rates less than 0.5 Hz were discarded. The mean firing rate for each cell was calculated as the mean number of spikes per second for the analysis recording section. To calculate the coefficient of variation (CV) of spiking activity for each cell, the inter-event interval histogram was extracted from the analysis recording section, and both the mean and the standard deviation was calculated. The CV was then calculated as the standard deviation divided by the mean IEI. Clustering analysis was performed on baseline bursting, frequency, and CV data using R (version 4.3.1) with R Studio. The R cluster package was used to calculate the dissimilarity matrix and perform hierarchical clustering; the R clValid package was used to perform internal cluster validation.

For grouped data, a Kolmogorov-Smirnov test was used to test normality. If one or more groups failed normality, non-parametric analyses were chosen. Two-tailed paired t-tests, one-way ANOVAs, or one-tailed t-tests were used when appropriate and stated. For analysis of changes in cell proportions, a Chi-squared test was used with numbers that were rounded to the nearest whole integer. All data are presented as mean ± SEM. Results were considered significant at an alpha of 0.05. Trends of p < 0.1 are reported.

### Blinding and randomization

Whenever possible, experimenters were blinded to treatment conditions, especially for stress experiments. Mice were assigned to control or stress conditions randomly. Furthermore, ChIs were assigned a CRF concentration randomly to ensure that we were not biasing our results based on baseline firing rate. As a result, firing rates across a particular group were not significantly different for the CRF concentration range.

## Results

### NAc shell ChIs have four distinct firing modes

We sampled a total of 145 ChIs (Cre+) from male and female *ChAT-ires-CRE^+/-^;Ai14 tdTomato* mice. Based on Bennet and Wilson’s observations of regular, irregular, and rhythmic bursting cell types(Bennett and Wilson, 1999), we classified cells first by observing whether or not the cell displayed bursting activity, defined as at least three burst events consisting of at least three spikes in a burst within a three-minute analysis period. Next, we used the firing rate of the cell and the coefficient of variation (CV) of the cell’s inter-event interval distribution, a measure of spike timing variability, to further determine the type of spiking pattern. Based on Bennet and Wilson’s as well as our own observations that firing rate tends to be negatively correlated with CV, we determined that using both metrics provides a more robust assessment of the cell’s spiking signature. Non-bursting cells with regular, single-spiking firing modes and narrow IEI distributions tended to have low CVs of 0.25 or less. Non-bursting irregular cells were characterized by irregularly-occurring single spikes, wider IEI distributions, and higher spike timing variability as indicated by CV values of 0.30 to 0.65. In contrast, rhythmic bursting cells displayed high-frequency bursts of activity followed either by complete pauses in activity or occasional single spikes; these cells usually had skewed or bimodal IEI distributions and the highest CV values typically of 0.70 to 1.00. However, in addition to these three cell types, we also observed some single spiking cells that displayed different degrees of spike timing variability as well as intermittent burst activity, with bursting patterns that tended to be longer in duration than bursts in rhythmic bursting cells. Since this cell spiking signature appeared to be a mix of both single spiking and bursting activity, we designated these as “mixed mode” ChIs. Figure 1a,b show example traces and corresponding IEI frequency histograms of these four different ChI firing patterns. Similar to Bennett and Wilson (1999), we observed a significant negative correlation between the firing rate and the IEI CV (r = -0.437, p < 0.0001), indicating that higher firing rates were associated with lower spike timing variability (Figure 1c). Firing rates tended to range between 1 and 4 Hz, but even within there were distinct and significant differences across the four ChI types (regular: 3.9±0.4 Hz; rhythmic: 2.3 ± 0.2 Hz; irregular: 1.8± 0.1 Hz; mixed mode: 3.3± 0.2 Hz, one-way ANOVA, F_3,144_ = 24.75, p < 0.0001, n = 145, see Figure 1d for Tukey’s post-hoc t-tests). Regular ChIs had significantly lower CVs compared to rhythmic or irregular firing patterns, while mixed mode ChIs had a significantly higher CV compared to regular ChIs (regular: 0.27± 0.01; rhythmic: 0.84 ± 0.07; irregular: 0.48 ± 0.02; mixed mode: 0.47± 0.03, Kruskal-Wallis test, p < 0.0001, n = 145, see Figure 1e for Dunn’s post-hoc comparison). Mixed mode neurons tended to have higher firing rates compared to irregular single spiking neurons, while having higher CVs compared to regular neurons due to the presence of bursts. Sorting ChIs based on our evaluation of burst activity, frequency, and CV, we observed that 47% of NAc shell ChIs displayed an irregular firing pattern, 27% had a regular firing pattern, 9% had a rhythmic bursting firing pattern and 17% had a mixed mode firing pattern (Figure 1f).

**Figure 1.**
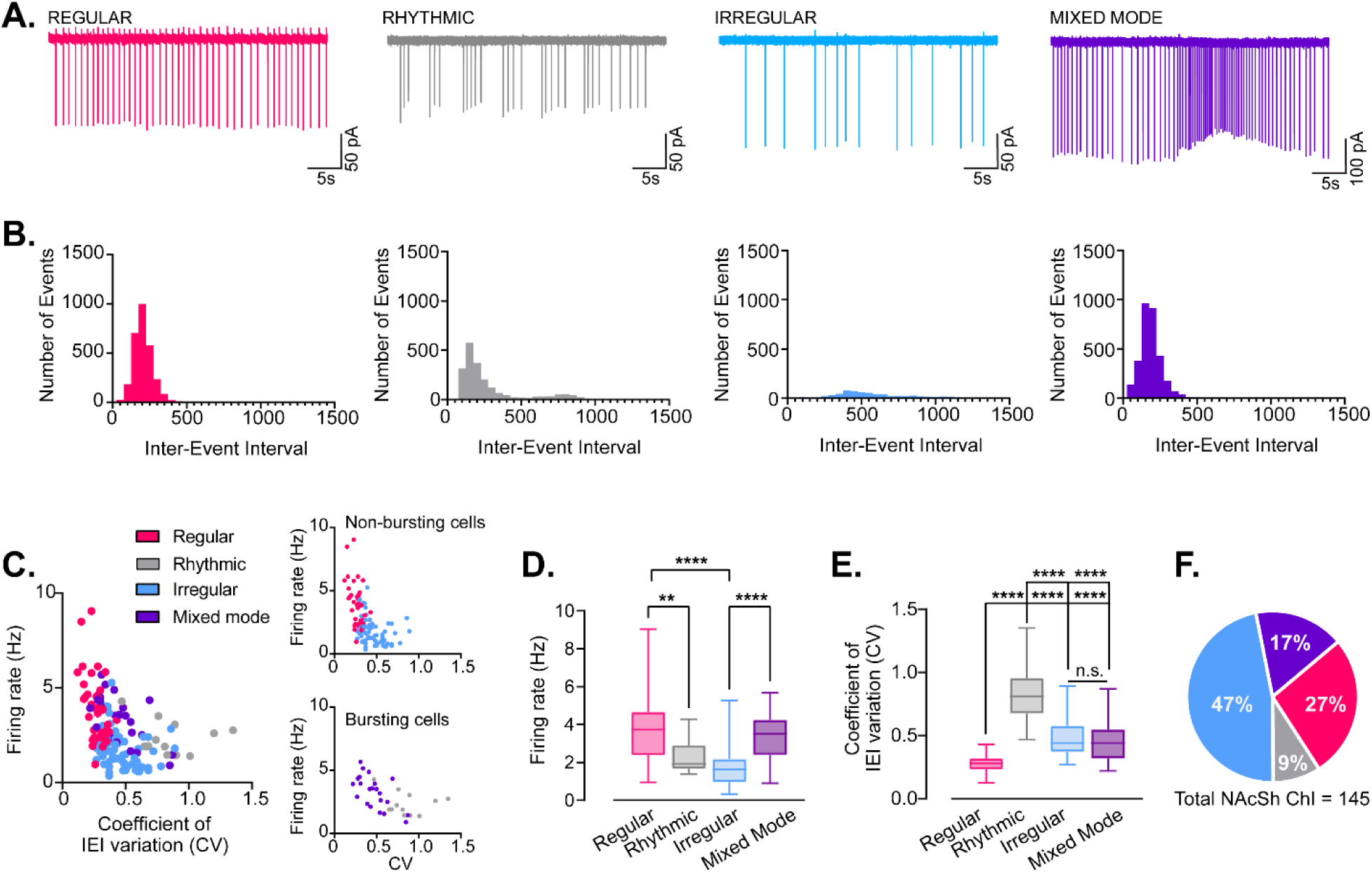
Cholinergic interneurons (ChIs) in the NAc shell have four distinct firing modes. A) Example traces of cell-attached recordings and B) corresponding frequency histograms of the inter-event interval made from regular, rhythmic, irregular and mixed mode ChIs recorded in the NAc shell. C) Firing rate x CV plot for 145 ChIs recorded in NAc shell. ChIs are color coded based on firing mode (regular = magenta, rhythmic = grey, irregular = light blue, mixed mode = purple). Insets separate the plot into non-bursting and bursting cells. There was significant negative correlation between firing rate and CV, r^2^ = -0.437, p < 0.0001. D) Summary data of firing rates for regular, rhythmic, irregular and mixed mode ChIs. One-way ANOVA with follow-up Tukey’s t-test. E) Summary data of CV for IEI for regular, rhythmic, irregular and mixed mode ChIs. Dunn’s post-hoc t-tests were used following a Kruskal-Wallis test. F) Pie chart of distribution of ChI spontaneous firing pattern from 145 cells recorded in NAc shell of male and female mice. *: Tukey’s or Dunn’s post-hoc t-tests, p < 0.05, **: Tukey’s or Dunn’s post-hoc t-test, p < 0.01, ***: Tukey’s or Dunn’s post-hoc t-test, p < 0.0001, ****: Tukey’s or Dunn’s post-hoc t-test, p < 0.00001

**Figure 2.**
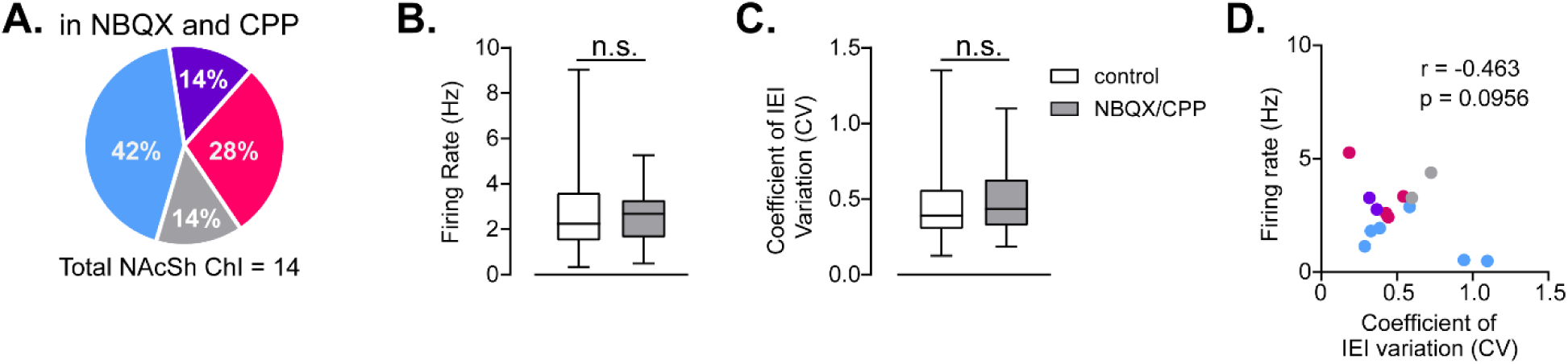
Blockade of glutamatergic synaptic transmission does not alter ChI firing properties. A) Pie chart of distribution of ChI spontaneous firing pattern from 14 recorded in NAc shell of male and female mice in the presence of NBQX and CPP. B) Summary data of firing rates for ChIs recorded in control ACSF (145) and ChIs recorded in the presence of NBQX and CPP (14). C) Summary data of CV of IEI for ChIs recorded in control ACSF (145) and ChIs recorded in the presence of NBQX and CPP (14). E) Firing rate x CV plot for 14 ChIs recorded in NAc shell in the presence of NBQX and CPP. ChIs are color coded based on firing mode (regular = magenta, rhythmic = grey, irregular = light blue, mixed mode = purple). There was a negative correlation between firing rate and CV, r^2^ = -0.463 that trended toward significance, p = 0.096. (regular = magenta, irregular = light blue, rhythmic bursting = grey, mixed mode = purple)

While this manual evaluation approach relied on both observing each cell’s spiking activity as well as utilizing burst, frequency, and CV metrics to identify ChI spiking types, an unbiased clustering approach was used to validate this method and determine whether ChIs could be naturally grouped based on these spiking metrics. Because quantifying bursting activity is a non-trivial challenge that varies highly across cell type and brain region and is particularly difficult in cells with varying spiking signatures, rather than quantifying burst features we instead chose to incorporate bursting as a categorical variable (0 = “Non-bursting”; 1 = “Bursting”) based on whether a cell showed at least three burst events within a thee-minute analysis period. We utilized hierarchical clustering since this approach allows us to easily visualize the relationships between clusters and to assess whether our observations of cell types and manual evaluation method shows concordance with this clustering algorithm. We first calculated the dissimilarity matrix, an essential first step in clustering since dissimilarity describes how different, or dissimilar, data points are in space. We used the *daisy* function in the R cluster package to calculate the Gower distance, a distance metric appropriate for scaling mixed variable types, including continuous and categorical (Figure S1a,b)(van de Velden, 2018). We next performed agglomerative hierarchical clustering on the Gower distances using the *hclust* function in the R cluster package. Ward’s linkage method provided the best agglomerative clustering coefficient (0.988 out of 1.000).

Adding labels representing our manual evaluation of each cell type to the resulting dendrogram revealed two clear clusters consisting of bursting and non-bursting cell types; bursting cell types were further divided into two groups mainly consisting of rhythmic bursting and mixed-mode cell types, and non-bursting cell types were divided into mostly regular and irregular cell types. These clustering results had a high degree of concordance with our observation of four main cell spiking types divided by bursting activity, suggesting that this is an appropriate grouping approach (Figure S1a). Lastly, we used an internal clustering validation method to assess the clustering structure of the hierarchical clustering result and compare it against other commonly used clustering methods, including position around medoids (PAM) and k-means. Using the clValid R package, we calculated two internal cluster validation measures, the Dunn index and the silhouette coefficient. These measures were calculated for hierarchical, PAM, and k-means clustering methods and across cluster solutions of 2-6 possible clusters. We found that hierarchical clustering resulting in two clusters provided the best structure, with a Dunn index of 1.023 and a silhouette coefficient of 0.785 (Figure S1c,d). While this validation measure confirmed that two groups provide the most straightforward clustering solution based on bursting activity, it is apparent from the hierarchical clustering results that these groups can be further divided into regular and irregular and rhythmic bursting vs mixed mode. This result provides support for the four distinct signatures of spiking activity we observed in the NAc and indicates that our evaluation approach provides a good amount of agreement with an unbiased clustering approach.*Effect of glutamatergic synaptic transmission on ChI spontaneous firing patterns*

Bennett and Wilson (1999) originally demonstrated that the heterogeneous ChI firing patterns were not driven by glutamatergic synaptic transmission. To confirm that this was also true for the NAc shell, we recorded from ChIs in the presence of antagonists for AMPA (NBQX, 5 µM) and NMDA (CPP, 5 µM) in male and female mice. The total proportions of regular, rhythmic, irregular, and mixed mode ChIs were preserved in the presence of glutamatergic synaptic blockers, even with a much smaller sample size (Chi-squared test, p = 0.6519, Figure 3a). There were no significant differences in the firing rate or CV in ChIs recorded in standard ACSF versus with the inclusion of AMPA and NMDA antagonists (control (n = 108) vs. NBQX/CPP (n = 14) - firing rate: 2.7± 0.2 Hz vs. 2.6 ± 0.4 Hz, Mann-Whitney test, p = 0.7947; CV: 0.48 ± 0.02 vs. 0.52 ± 0.07, Mann-Whitney test, p = 0.6633, Figure 3b,c). Finally, though not statistically significant, there was a qualitatively similar negative correlation between firing rate and CV as seen in control ACSF (r = -0.463, p = 0.096). These data are consistent with previous findings in the dorsal striatum that spontaneous firing patterns are not driven by glutamatergic synaptic transmission.

**Figure 3.**
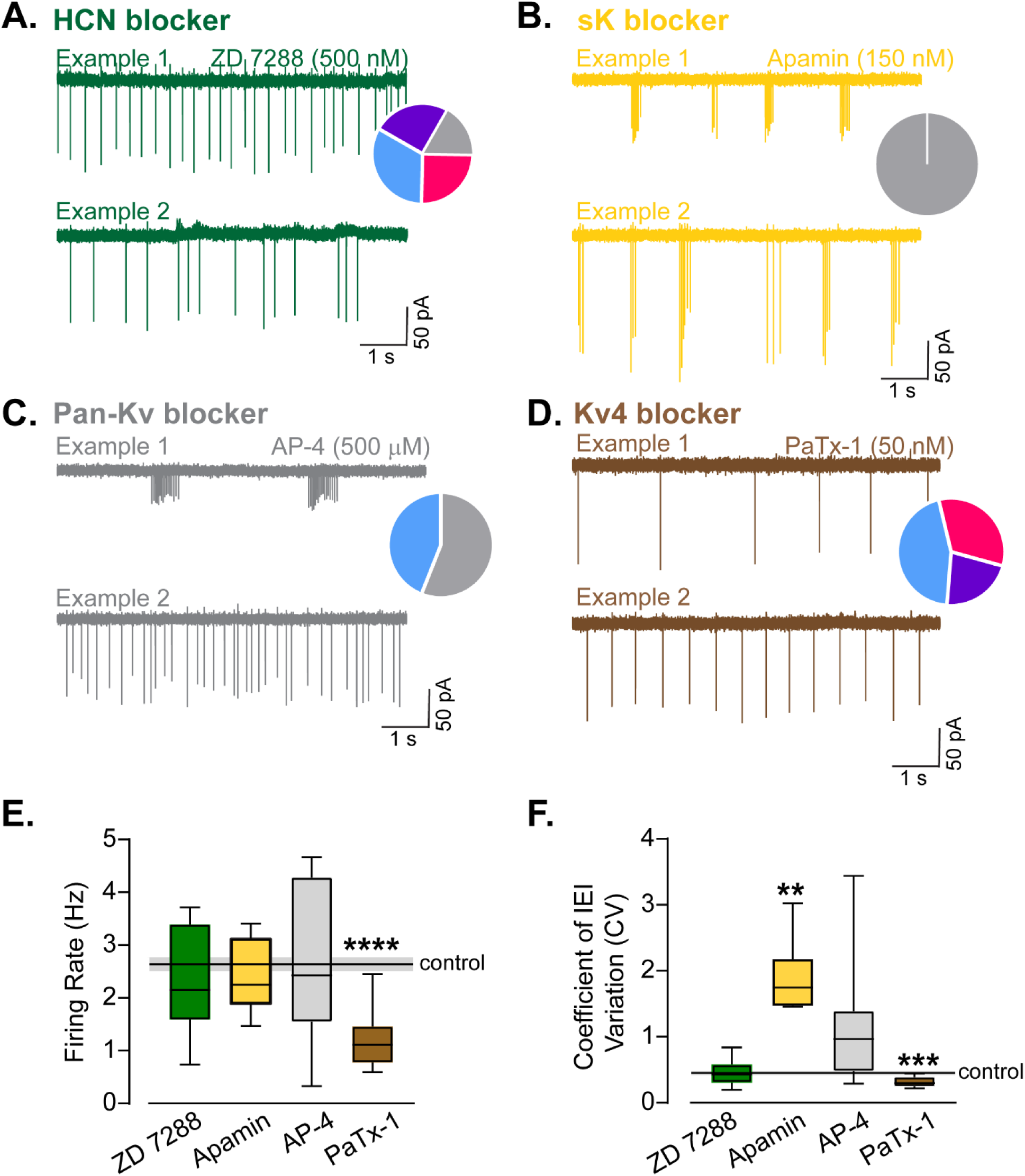
Effect of ion channel blockers on ChI firing patterns. A-D) Example traces from cell-attached recordings made from ChIs in the NAc shell in the presence of the HCN blocker, ZD 7288 (500 nM) (A), sK blocker, Apamin (150 nM) (B), Pan-Kv blocker, AP-4 (500 µM)(C) and Kv4.2 blocker, Phrixotoxin-1 (50 nm)(D). Insets represent the firing pattern distribution observed across ChIs for each of the drug conditions (regular = magenta, irregular = light blue, rhythmic bursting = grey, mixed mode = purple). E) Mean firing rate of ChIs recorded in the different drug conditions compared to the average firing rate of ChIs recorded in drug free ACSF (denoted by solid black line, with standard error around the mean in grey)(**** = one sample t-test, p < 0.00001). F) Mean CV of IEI of ChIs recorded in the different drug conditions compared to the average CV of IEI of ChIs recorded in drug free ACSF (denoted by solid black line, with standard error around the mean in grey)( one sample t-tests, ** = p < 0.01, *** = p < 0.001).

### Impact of ion channel blockers on ChI firing patterns in the NAc shell

We next investigated changes in firing rate, CV and firing pattern distributions following the attenuation of different ion channels known to regulate firing patterns generally and ChIs specifically (Bennett, Callaway and Wilson, 2000; Cheng et al., 2019; Oswald et al., 2009; Song et al., 1998). We compared our results to the mean ChI firing rate, CV and firing pattern distribution of cells recorded in ACSF (Figure 3). Attenuation of the hyperpolarization cyclic nucleotide gated (HCN) cation channel current (I_H_) using ZD 7288 produced little change in the mean firing rate and CV compared to the average of all ChIs recorded in ACSF (one-sample t-tests, n = 12), but did seem to increase in the overall heterogeneity (Chi-squared test, p = 0.0024) (Figure 3a,e-f). Attenuation of sK channels with apamin led to a significant shift in the proportions of firing patterns toward entirely rhythmic firing activity (Chi-squared test, p < 0.0001)(Figure 3b, e-f), indicated by a dramatically altered ChI CV of the IEI (one-sample t-test, p = 0.0021), with no change in the overall spiking frequency (one-sample t-test, p = 0.4513, n = 6).

Attenuation of voltage-activated potassium channels had mixed effects that led to either irregular or highly rhythmic firing patterns that averaged to no change in mean firing rate or CV (one-sample t-tests, n = 12), but again produced a significant change in the proportions of firing patterns (Chi-squared, p < 0.0001). Kv4.2 channels that mediate an A-type K+ current have been shown to drive irregular firing in the cortex and hypothalamus (Mendonca et al., 2016; Mendonca et al., 2018). Furthermore, Kv 4.2 has been shown to be highly expressed in ChIs, and depolarization-activated somatodendritic K+ currents in cholinergic interneurons are dominated by Kv4.2 and Kv4.1(Song et al., 1998). Thus, we investigated whether blocking Kv4.2 with Phrixotoxin-1 (PaTx-1) would reduce the proportions of irregular or rhythmic firing neurons. We found that PaTx-1 (50 nM) reduced the firing rate (one-sample t-test, p < 0.0001, n = 9) and the CV (one-sample-test, p = 0.0002), indicating a shift toward more regular firing. While cells retained their irregular patterns, rhythmic firing cells were completely absent, producing a significant shift in firing pattern proportions compared to control (Chi-squared test, p < 0.0001)

### Differential CRF sensitivity based on ChI spontaneous firing patterns

CRF is released in the NAc in response to salient stimuli and potentiates ChI firing via sK channels (Lemos, Shin and Alvarez, 2019; Lemos et al., 2012). Given the importance of sK channels in determining dorsal striatal ChI firing properties (Bennett, Callaway and Wilson, 2000; Bennett and Wilson, 1999) and our findings in the NAc shell that blockage of sK shifts activity away from regular spiking toward rhythmic bursting (Figure 3), we speculated that CRF may be a reasonable molecular candidate for how salient stimuli alter the firing properties of ChIs in the NAc. We assessed the ability of CRF to alter the firing rate, CV, and overall firing pattern of NAc shell ChIs collected from male and female mice, segregated into the four classifications based on their baseline firing patterns. We chose concentrations based on previous work done in the NAc core (Lemos, Shin and Alvarez, 2019). At the maximal concentration (100 nM), CRF significantly increased the firing rate of all four categories of ChI firing patterns (one-sample t-tests compared to 100% baseline, n = 9-20, Figure 4a,b,d,e,h,i,l,k).

**Figure 4.**
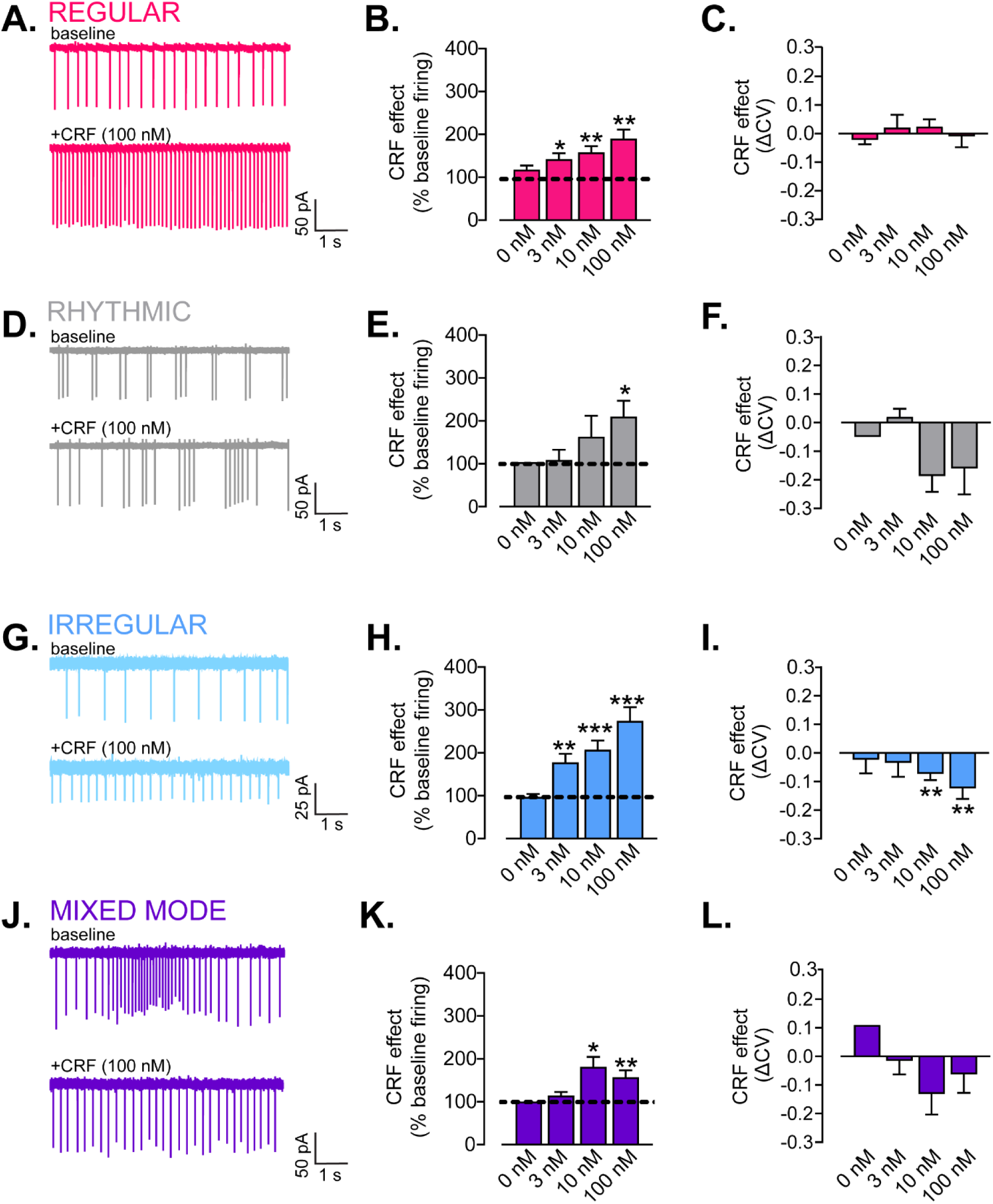
Differential sensitivity to CRF based on ChI firing patterns. A) Example trace of cell-attached recordings prior to and following CRF in from a Regular ChI. B) Summary data of CRF effect on firing rate (% baseline) for 0, 3, 10, 100 nM in Regular ChIs. C) Summary data of CRF effect on CV of IEI (ΔCV) for 0, 3, 10, 100 nM in Regular ChIs. D) Example trace of cell-attached recordings prior to and following CRF in from a Rhythmic ChI. E) Summary data of CRF effect on firing rate (% baseline) for 0, 3, 10, 100 nM in Rhytmic ChIs. F) Summary data of CRF effect on CV of IEI (ΔCV) for 0, 3, 10, 100 nM in Rhythmic ChIs. G) Example trace of cell-attached recordings prior to and following CRF in from a Irregular ChI. H) Summary data of CRF effect on firing rate (% baseline) for 0, 3, 10, 100 nM in Irregular ChIs. I) Summary data of CRF effect on CV of IEI (ΔCV) for 0, 3, 10, 100 nM in Irregular ChIs. J) Example trace of cell-attached recordings prior to and following CRF in from a Mixed Mode ChI. K) Summary data of CRF effect on firing rate (% baseline) for 0, 3, 10, 100 nM in Mixed Mode ChIs. L) Summary data of CRF effect on CV of IEI (ΔCV) for 0, 3, 10, 100 nM in Mixed Mode ChIs. *: One-sample-test vs. 100%, p < 0.05,**: One-sample-test vs. 100%, p < 0.01,***: One-sample-test vs. 100%, p < 0.0001,*****: One-sample-test vs. 100%,, p < 0.0001

However, for 3 nM and 10 nM CRF, the responses varied depending on the basal spontaneous firing pattern of ChIs. CRF (10 nM) increased the firing rate above baseline significantly for regular, irregular, and mixed mode ChIs but not for rhythmic ChIs (one-sample t-tests compared to 100% baseline, n = 3-24, Figure 4b,e,I,k). CRF (3 nM) only increased the firing rate above baseline significantly for regular and irregular ChIs (one-sample t-tests compared to 100% baseline, n = 10-24, Figure 4b,h). CRF (10, 100 nM) significantly reduced the IEI CV in irregular ChIs only, with no significant effect on regular, rhythmic, or mixed mode ChIs (one-sample t-tests compared to 0, n = 3-24, Figure 4c,g,j,m).

We next directly compared the change in firing rate induced by 3, 10 and 100 nM CRF based on firing pattern. There was a trend for differences in response at the low concentration (one-way ANOVA, F_3,31_ = 2.354, p = 0.0912, n = 3-15; Figure 5a). There were no significant differences at the 10 nM concentration (Figure 5b). In contrast, at the maximal concentration, CRF differentially potentiated the ChI firing rate based on the basal spontaneous firing pattern (one-way ANOVA, F_3,48_ = 3.506, p = 0.0222, Tukey’s post-hoc t-test, mixed mode vs irregular, p = 0.0306, n = 3-20, Figure 5c). We had previously shown that there was a negative correlation between CRF response and baseline firing rate (Lemos, Shin and Alvarez, 2019). We replicated this finding, demonstrating that the CRF response (percent of baseline firing) is negatively correlated with the baseline firing rate, best fit with a semi-log line (r^2^ = -0.60, p < 0.001, Figure 5d). Finally, we assessed the firing patterns prior to and following either vehicle or CRF application (100 nM). Similar to the ChI firing pattern distribution assessed from all 145 cells, at baseline, the ChIs were composed of 33% regular, 7% rhythmic, 53% irregular and 7% mixed mode patterns before vehicle application. Before CRF application (100 nM), ChIs were composed of 24% regular, 9% rhythmic, 50% irregular and 17% mixed mode patterns. There was no significant effect of vehicle application during the same timecourse as CRF application, indicating that the experimental paradigm in and of itself is not causing artefactual shifts in firing patterns (Chi-squared, p = 0.4936, n = 15, Figure 5e). Following CRF (100 nM), these proportions were shifted to 42% regular, 5% rhythmic, 30% irregular and 23% mixed mode (Chi-squared test, p = 0.0011, n = 52, Figure 5f).

**Figure 5.**
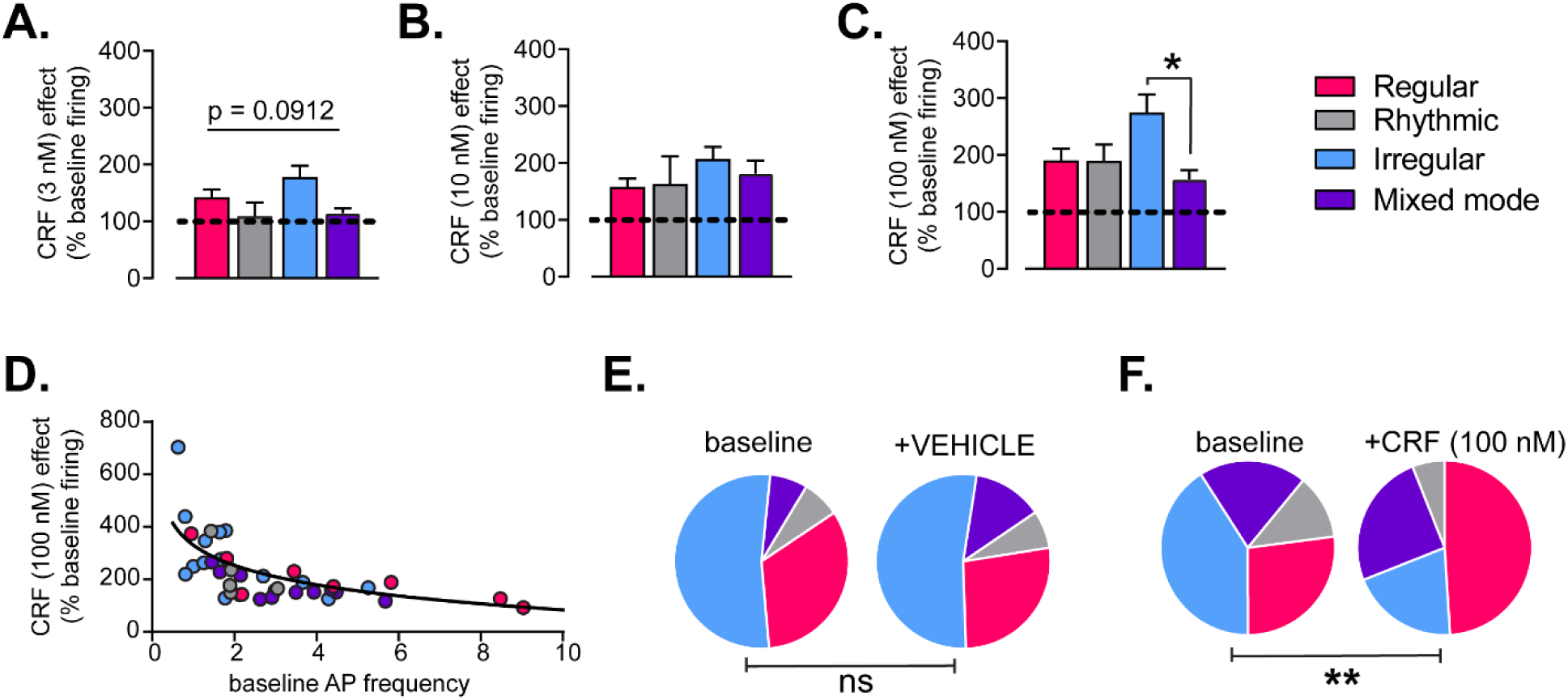
CRF shifts ChI firing properties toward a regular firing pattern. A) Summary data of CRF (3 nM) effect on firing rate (% baseline) for regular, rhythmic, irregular and mixed mode ChIs. B) Summary data of CRF (10 nM) effect on firing rate (% baseline) for regular, rhythmic, irregular and mixed mode ChIs. C) Summary data of CRF (100 nM) effect on firing rate (% baseline) for regular, rhythmic, irregular and mixed mode ChIs. One-way ANOVAs, *: Tukey’s post-hoc t-tests, p < 0.05, **: Tukey’s post-hoc t-test, p < 0.01, ***: Tukey’s post-hoc t-test, p < 0.0001, ****: Tukey’s post-hoc t-test, p < 0.00001. D) CRF 100 nM response (%baseline) x baseline firing rate plot for 39 ChIs recorded in NAc shell prior to and following CRF. ChIs are color coded based on firing mode (regular = magenta, rhythmic = grey, irregular = light blue, mixed mode = purple). Data is fit with a semi-log line, r^2^ = -0.60, p < 0.001. E) Firing pattern distribution in ChIs prior to (left) and following (right) vehicle bath application. n.s. = Chi-squared test. F) Firing pattern distribution in ChIs prior to (left) and following (right) CRF (100 nM) bath application. ** = Chi-squared test, p < 0.01.

### Sex differences in the basal functional properties of NAc shell ChIs

To our knowledge, sex-dependent differences in ChI properties have not been assessed. We analyzed differences in firing rates, IEI CVs, and firing mode proportions between male and female mice, and in females across the estrous cycle. On average, males had significantly higher firing rates than females (male: 3.4± 0.3 Hz, n = 30; female 2.5 ± 0.2 Hz, n = 116, Mann-Whitney test, p =0.0089, Figure 6a,b). As described in the methods, we tracked the female estrous cycle over an 8–10-day period to be confident of estrous cycle stage on the day of sacrifice. A final vaginal lavage was performed prior to sacrifice, and the cycle stage was visualized and noted (Figure 6d). While firing rates in females appeared lower than males across the estrous cycle, this was only significant in estrus females when the female data was parsed (male: 3.4± 0.4 Hz, n = 30; female-proestrus: 2.5 ± 0.4 Hz, n = 24; female-estrus: 2.2 ± 0.2 Hz, n = 32; female-metestrus: 2.6 ± 0.3 Hz, n = 20, female-diestrus: 2.7 ± 0.3 Hz, n = 37, Kruskal-Wallis test, p = 0.0589, Dunnett’s comparison test, male vs. female-estrus, p = 0.0286, Figure 2e). Notably, the variance across groups for both firing rates and CVs were significantly different (Bartlett’s test, p = 0.0112). On average, males and females did not have significantly different IEI CVs (male: 0.44 ± 0.03, n = 30; female 0.45 ± 0.02, n = 116, Mann-Whitney, p =0.5357, Figure 6c). Likewise, when comparing the CV of male and females across the estrous cycle, no significant differences were found (Kruskal-Wallis test, p = 0.1435, Figure 6f). We next sorted our previously classified cells by sex and by estrous cycle. Interestingly, we found a significant difference in the proportion of ChI firing modes between males and females, driven by a higher proportional of “mixed mode” neurons and a lower proportion of “regular” neurons in female mice (Chi-square test, p = 0.0244, Figure 6g). There were also robust differences in cell type proportions across the estrous cycle (Chi-square test, p = 0.0134, Figure 6h). Notably, ChIs collected in the estrus phase had higher proportions of mixed mode and rhythmic bursting firing patterns. ChIs from diestrus females had a very high proportion of irregular firing neurons with very low proportions of regular, mixed mode and rhythmic firing patterns, compared to ChIs collected in other cycle stages. Taken together, the data demonstrate differences in ChI firing rates and patterns between males and females.

**Figure 6.**
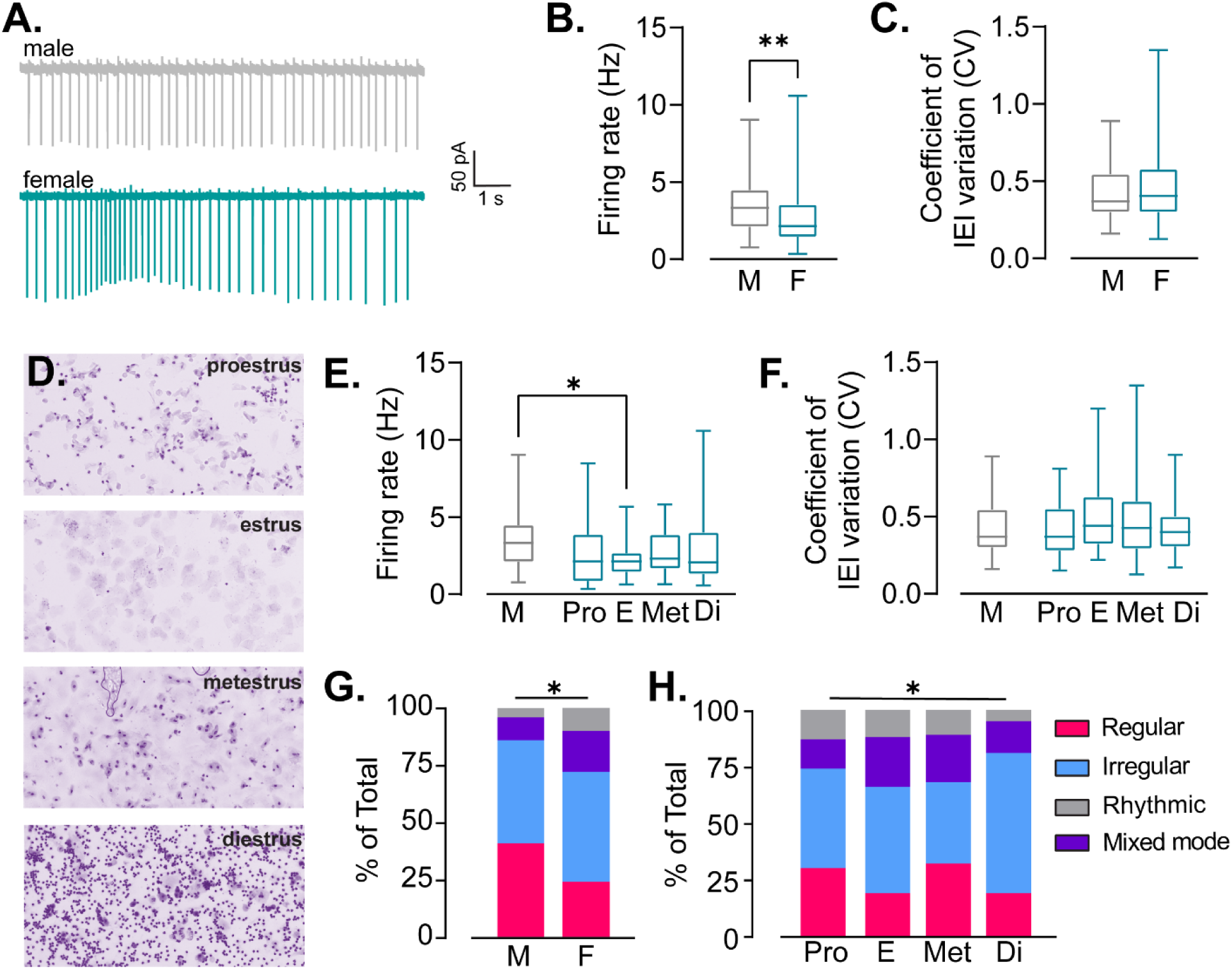
Sex differences in NAc shell ChI firing properties. A) Example traces of cell-attached recordings made from NAc shell ChIs in male (top) and female (bottom) mice. B) Summary data of firing rates recorded in male and female mice. Mann-Whitney test, ** p < 0.01. C) Summary data of CV of IEI for ChIs recorded in male and female mice. E) Example images of vaginal cytology used to classify females into proestrus, estrus, metestrus and diestrus cycle stages. E) Summary data of firing rates recorded in NAc shell ChI from male and female mice across the estrous cycle. Dunn’s post-hoc t-tests were used following a Kruskal-Wallis test: * = Male vs. Estrus, p< 0.05. F) Summary data of CV of IEI recorded in NAc shell ChI from male and female mice across the estrous cycle. G) Distribution of ChI firing mode in male and females (pooled, left) across estrous cycle (separated, right). * = Chi-squared test of male vs. female,p < 0.05; Chi-squared test of female across estrous cycle, p < 0.05.

### Difference in CRF sensitivity across the estrous cycle

Since there were significant difference in firing rate and patterns between males and females, and between females during different cycles of estrous, we hypothesized that there would be differences in CRF sensitivity given our finding that CRF sensitivity was different depending on the firing pattern. Overall, we did not observe any differences between males and pooled females in the CRF potentiation of ChI firing or the change in CV (Figure 7a,b). We also did not observe any significant differences in CRF’s impact on the CV across the estrous cycle (data not shown). However, as we would expect given the higher proportion of irregular firing neurons, we did find that a low concentration of CRF (3 nM) potentiated ChI firing rate in diestrus females compared to females in estrus and to less extent in metestrus (timecourse: Mixed-effects model, REML, p > 0.05, Fixed effects Type III, significant effect of time, p < 0.0001, trend for effect of estrous, p = 0.0988; one-way ANOVA on the maximal effect, F_2,20_ = 4.3539, p = 0.0237; Tukey’s post-hoc t-test, diestrus vs estrus, p = 0.0473, diestrus vs metestrus, p = 0.0565, n = 6-10, Figure 7c,d). Proestrus is a very challenging stage of the estrous cycle to capture due to its short duration compared to other stages. As a result, we were not able to collect enough proestrus samples at this concentration, and the cells we did collect that were assigned to 3 nM were all very low firing cells (0.5-1 Hz), making this data uninterpretable. We therefore excluded proestrus from this analysis. We did not find significant differences in CRF sensitivity across the estrous cycle for the 10 nM or 100 nM CRF concentrations (n = 8-12 for 10 nM and n = 7-12 for 100 nM; analyses run as above, p-values > 0.05, Figure 7e-h).

**Figure 7.**
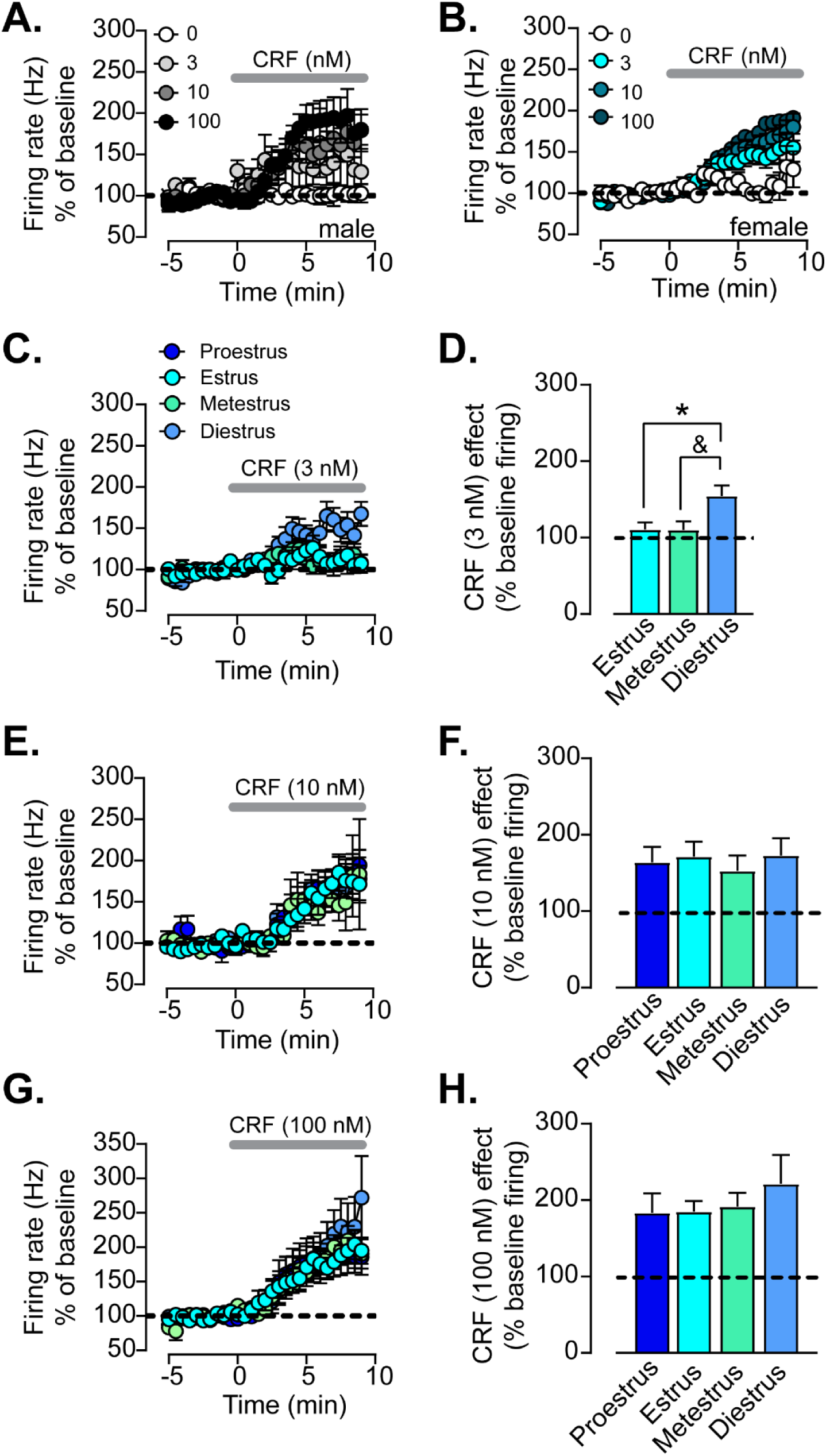
Estrous cycle dependent difference in sensitivity to a low concentration of CRF. A) Timecourse of normalized firing rate (% baseline) prior to and following bath application of range of concentrations of CRF (0, 3, 10, 100 nM, collected in separate slices) in coronal sections of NAc shell from male mice. B) Timecourse of normalized firing rate (% baseline) prior to and following bath application of range of concentrations of CRF (0, 3, 10, 100 nM, collected in separate slices) in coronal sections of NAc shell from female mice. C) Timecourse of normalized firing rate (% baseline) prior to and following bath application of 3 nM CRF from females in estrus, metestrus and diestrus. D) Average maximal CRF (3nM) response (% baseline) in females in estrus, metestrus and diestrus. One-way ANOVA, * = Tukey’s post-hoc t-test, p < 0.05, & = Tukey’s post-hoc t-test, p = 0.0565). E) Timecourse of normalized firing rate (% baseline) prior to and following bath application of 10 nM CRF from females in proestrus, estrus, metestrus and diestrus. F) Average maximal CRF (10 nM) response (% baseline) in females in proestrus, estrus, metestrus and diestrus. G) Timecourse of normalized firing rate (% baseline) prior to and following bath application of 100 nM CRF from females in proestrus, estrus, metestrus and diestrus. H) Average maximal CRF (100 nM) response (% baseline) in females in proestrus, estrus, metestrus and diestrus.

### Repeated stressor exposure impacts Crh and Crhr1 mRNA levels

We next sought to determine whether a behavioral treatment that potentially elevates CRF levels in the NAc, possibly emulating CRF bath application, would shift the firing pattern distribution of ChIs toward a larger proportion of regular firing ChIs. While it is very difficult to assess CRF release directly, there is a group of CRF-containing neurons within the NAc that may release CRF onto surrounding ChIs (Itoga et al., 2019). Importantly, we can use reliable RNAScope methodologies to assess changes in *Crh* mRNA levels within the NAc following repeated swim stress compared to control animals. We killed male mice and flash froze their brains 7 days after the final swim stress exposure. For these stress experiments as well as the ones described below, we chose to use males since we had just established females have varying ChI firing properties and patterns across baseline as well as varying CRF sensitivity in the stress-naïve state. We found that *Crh* mRNA total puncta was higher in the NAc of stress-exposed animals compared to unperturbed animals (control: 106.7 ± 5.8; FSS: 143.1 ± 15.95, t-test, p = 0.03, n = 44-46 unique images from 3 animals each, Figure 8a,b). This did not translate into a significant change in the percent of *Crh*+ cells in the NAc (unpaired t-test, p = 0.3655, n = 3 animals each, Figure 8c). We also quantified the number of CRF-R1+ CINs and puncta per ChI from the same animals by multiplexing for *Chat and Crhr1* mRNA. We found no significant differences in the number of puncta per CIN cell, though there was a trend for an increase in *Crhr1* puncta in stress-exposed mice, this was not signficant (naïve – 67 unique ChIs: 21 ± 1 puncta; stress – 71 unique ChIs: 25 ± 1.5, p = 0.0589, Figure 8d,f) or percent of CRF-R1+ ChIs (Chi-squared test, p > 0.05, n = 3 each, Figure 8e). CRF microinfusion into the NAc increases locomotor behavior, and novelty exposure triggers CRF release in the NAc(Holahan, Kalin and Kelley, 1997; Lemos et al., 2012). We have previously shown that females have an elevated locomotor response to a novel environment compared to males (Razidlo et al., 2022). Here, as proof of concept, we tested whether repeated swim stress, which led to elevations in NAc *Crh* levels, impacted novelty-induced locomotor behavior. We found that forced swim stress (FSS) reduced locomotor habituation to a novel environment compared to control conditions (two-way RM ANOVA, time x stress treatment interaction, F_1,8_ = 46.47, p = 0.0001, n = 5 each). This finding provides evidence that stress can alter NAc CRF-dependent behaviors.

**Figure 8.**
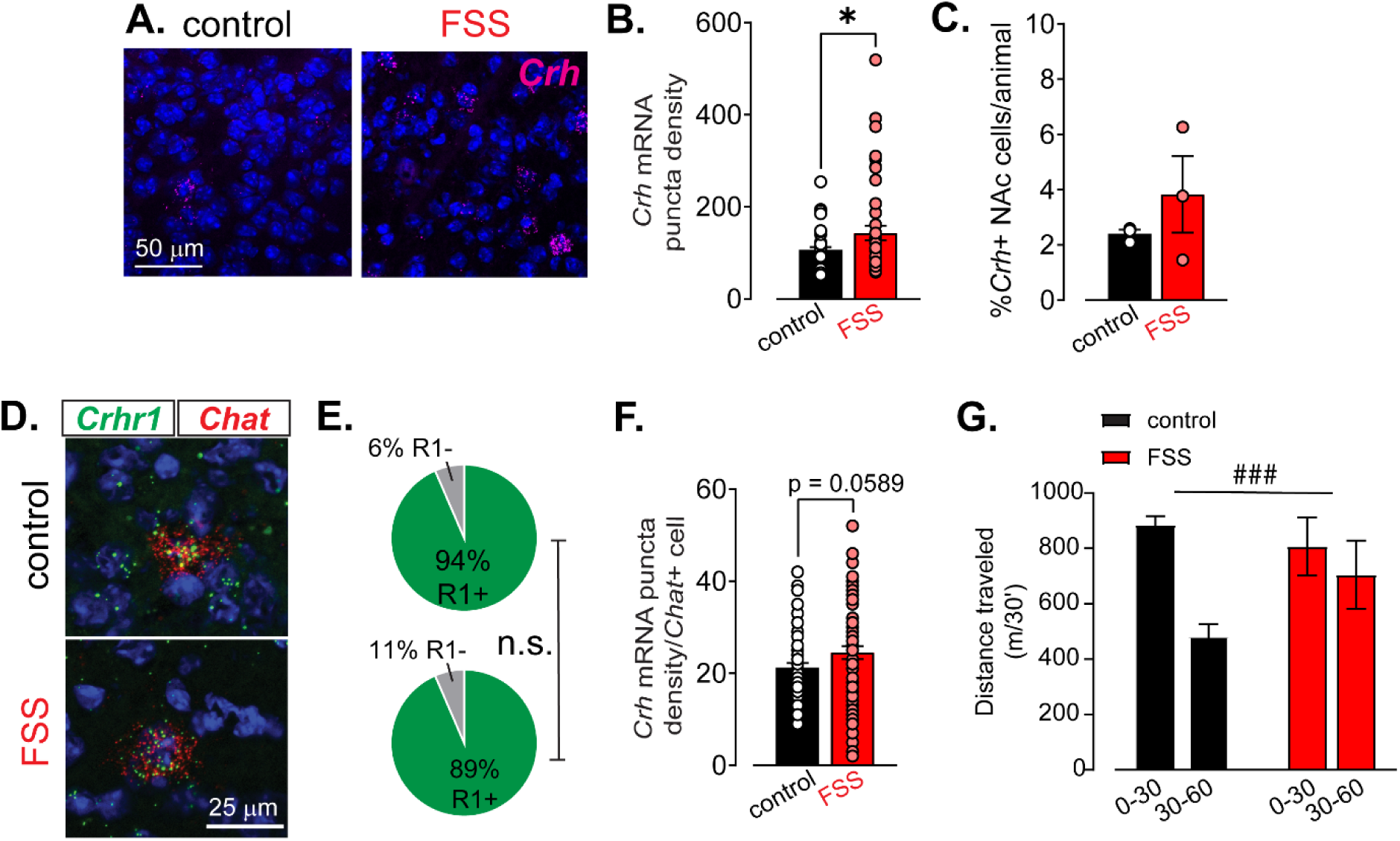
Repeated stressor exposure increases *Crh* mRNA levels in the NAc. A) Example images of *Crh* mRNA (magenta) with DAPI counterstain in NAc collected from a control male and FSS exposed male using in fluorescent situ hybridization techniques. B) Summary data of number of *Crh* mRNA puncta per DAPI cell collected across 3 animals per behavioral treatment. * = unpaired t-test, p < 0.05. C) Summary data of %*Crh* + cells in control and FSS exposed mice, n = 3 per group. D) Example images from fluorescent in situ hybridization experiments multiplexing for *Crhr1* mRNA (green) and *Chat* mRNA (red) with DAPI counterstain in NAc collected from a control male and FSS exposed male. E) Pie charts showing distribution of *Crhr1+/Chat+* and *Crhr1-/Chat+* cells in the NAc of control and stress-exposed male mice. n.s. = Chi-squared test, p > 0.05. F) Summary data of number of *Crhr1* mRNA puncta per *Chat+* cell collected across 3 animals per behavioral treatment, unpaired t-test, p = 0.0589. G) Distance traveled (m/30’) in the first 30 (0-30) and last 30 (30-60) of a 60-minute novel open field test conducted in control and FSS-exposed male mice. ### = Two-way ANOVA, time x stress treatment interaction, p < 0.001.

### Repeated stressor exposure shifts ChI firing distribution and CRF sensitivity in the NAc core but not the NAc shell

We next assessed the impact of stressor exposure on the firing properties and patterns in the NAc core and the NAc shell as well as assessed stress-induced changes in CRF sensitivity (3, 10, 100 nM). We chose to examine effects at approximately 1 (6-8 days) and 2 (14-18 days) weeks post-stressor exposure to determine the persistence of the effects. Again, we chose to use males for these experiments to avoid the additional complexity of estrous-driven changes shown above. Furthermore, we chose to use sagittal sections for these experiments since, in our experience, distinguishing core and shell across the medial-lateral axis is more reliable in sagittal sections compared to coronal sections. We confirmed that the plane of section did not significantly affect the firing rate, CV or firing patterns (Figure S2). We first noted striking differences in the proportions of ChI firing patterns between NAc core and NAc shell in control animals. While the firing rate was not different between shell and core regions, the NAc shell had a significantly higher IEI CV than NAc core (firing rate: NAc core control: 3.4 ± 0.3 Hz, NAc shell control: 3.4 ± 0.3 Hz, unpaired t-test, p = 0.8292; CV: NAc core: 0.23 ± 0.02, NAc shell control: 0.38 ± 0.03, unpaired t-test, p = 0.0005, n = 16-23, Figure S3). This was due to the predominance of regular firing neurons in the NAc core, with only a small percentages of mixed mode, irregular or rhythmic firing neurons.

Strikingly, repeated stressor exposure increased the IEI CV and increased the heterogeneity of ChI firing patterns recorded in the NAc core following 1 week of incubation. This was fully reversed two weeks post-stressor exposure. We did not observe a change in the firing rate (firing rate: control: 3.4 ± 0.3 Hz, FSS-1 week: 3.9 ± 0.7 Hz, FSS-2 week: 4.5 ± 0.6 Hz, one-way ANOVA, F_2,46_ = 0.9145, p = 0.4086; CV: control: 0.22 ± 0.02, FSS-1 week: 0.43 ± 0.06, FSS-2 week: 0.25 ± 0.03, one-way ANOVA, F_2,46_ = 5.682, p = 0.0062; pattern distribution: control: 81% regular, 6% irregular, 13% mixed mode, 0% rhythmic; FSS-1 week: 48% regular, 42% irregular, 5% mixed mode, 5% rhythmic; FSS-2 week: 72% regular, 14% irregular, 14% mixed mode, 0% rhythmic, Chi-squared tests, p-values < 0.0001, n = 14-19, Figure 9a-d).

**Figure 9.**
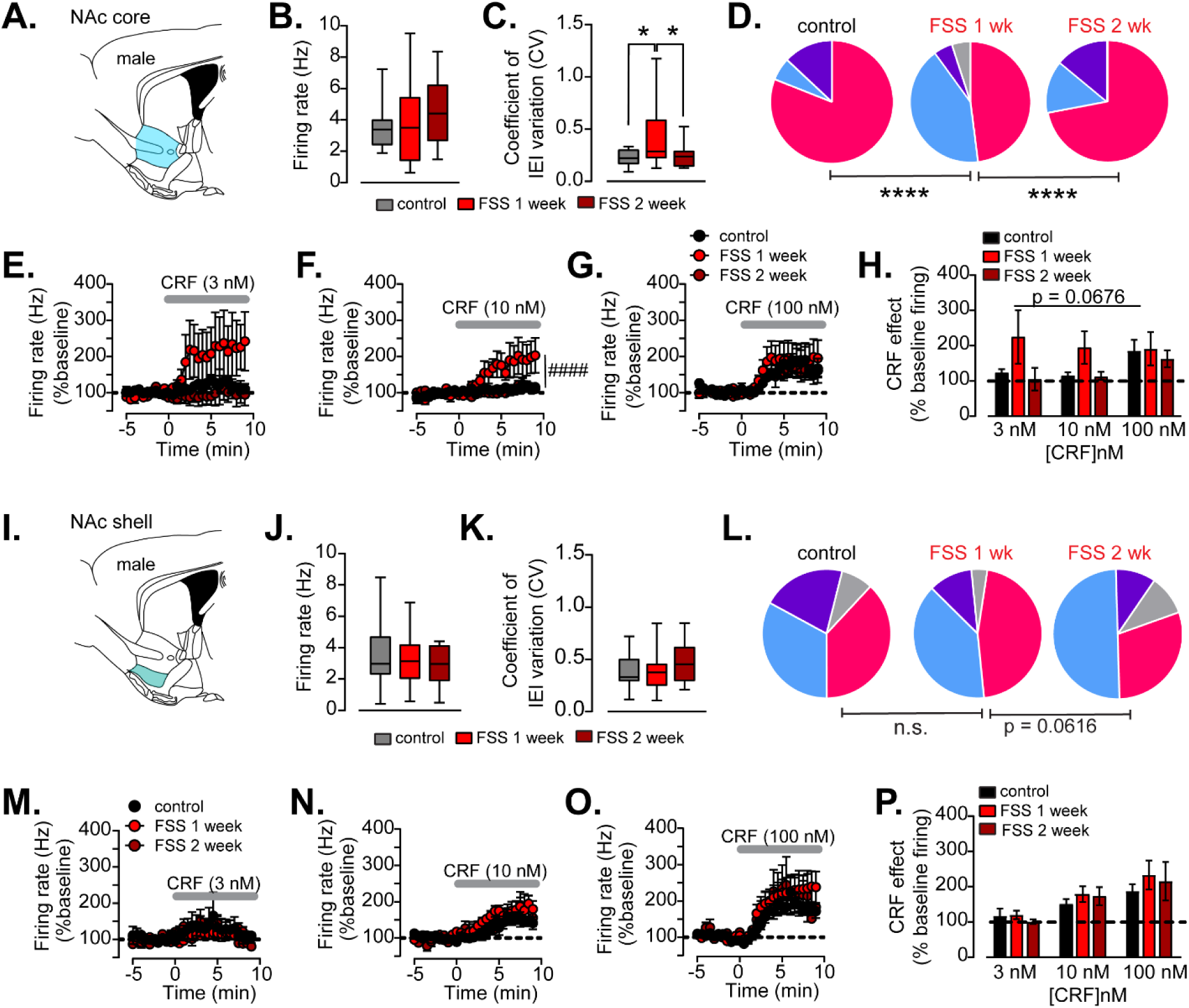
Repeated stressor exposure impacts ChI firing pattern distribution and CRF sensitivity in the NAc. A) Schematic of NAc core region in a sagittal section. B) Mean baseline firing rate of ChIs recorded from the NAc core of control, FSS-1 week incubation, FSS-2 week incubation male mice. C) Mean baseline CV of IEI of ChIs recorded from the NAc core of control, FSS-1 week incubation, FSS-2 week incubation male mice. One-way ANOVA, * = Tukey’s post-hoc t-tests, p-values < 0.05. D) Firing pattern distribution of ChIs recorded from the NAc core of control, FSS-1 week incubation, FSS-2 week incubation male mice. ****Chi-squared tests, p < 0.0001. (regular = magenta, irregular = light blue, rhythmic bursting = grey, mixed mode = purple). E) Timecourse of normalized firing rate (% baseline) prior to and following bath application of 3 nM CRF in ChIs recorded from the NAc core of control, FSS-1 week incubation, FSS-2 week incubation male mice. F) Timecourse of normalized firing rate (% baseline) prior to and following bath application of 10 nM CRF in ChIs recorded from the NAc core of control, FSS-1 week incubation, FSS-2 week incubation male mice, Two-Way RM ANOVA, #### = time x stress treatment interaction, p < 0.0001. G) Timecourse of normalized firing rate (% baseline) prior to and following bath application of 100 nM CRF in ChIs recorded from the NAc core of control, FSS-1 week incubation, FSS-2 week incubation male mice. H) Average maximal CRF (3,10,100 nM) response (% baseline) in NAc core of control, FSS-1 week incubation, FSS-2 week incubation male mice. I) Schematic of NAc shell region in a sagittal section. J) Mean baseline firing rate of ChIs recorded from the NAc shell of control, FSS-1 week incubation, FSS-2 week incubation male mice. K) Mean baseline CV of IEI of ChIs recorded from the NAc shell of control, FSS-1 week incubation, FSS-2 week incubation male mice. L) Firing pattern distribution of ChIs recorded from the NAc shell of control, FSS-1 week incubation, FSS-2 week incubation male mice. (regular = magenta, irregular = light blue, rhythmic bursting = grey, mixed mode = purple). M) Timecourse of normalized firing rate (% baseline) prior to and following bath application of 3 nM CRF in ChIs recorded from the NAc shell of control, FSS-1 week incubation, FSS-2 week incubation male mice. N) Timecourse of normalized firing rate (% baseline) prior to and following bath application of 10 nM CRF in ChIs recorded from the NAc shell of control, FSS-1 week incubation, FSS-2 week incubation male mice. O) Timecourse of normalized firing rate (% baseline) prior to and following bath application of 100 nM CRF in ChIs recorded from the NAc shell of control, FSS-1 week incubation, FSS-2 week incubation male mice. P) Average maximal CRF (3,10,100 nM) response (% baseline) NAc shell of control, FSS-1 week incubation, FSS-2 week incubation male mice.

In contrast, repeated stressor exposure had no significant effect on the firing rate, CV or firing pattern proportions in the NAc shell, disproving our initial hypothesis that stressor exposure may replicate the effects of acute CRF bath application (firing rate: control: 3.5 ± 0.4 Hz, FSS-1 week: 3.2 ± 0.3 Hz, FSS-2 week: 2.9 ± 0.4 Hz, one-way ANOVA, F_2,58_ = 0.6266, p = 0.5379; CV: control: 0.38 ± 0.03, FSS-1 week: 0.37 ± 0.03, FSS-2 week: 0.48 ± 0.07, one-way ANOVA, F_2,58_ = 1.632, p = 0.2045; pattern distribution: control: 38% regular, 33% irregular, 21% mixed mode, 8% rhythmic; FSS-1 week: 46% regular, 39% irregular, 11% mixed mode, 4% rhythmic; FSS-2 week: 30% regular, 50% irregular, 10% mixed mode,10% rhythmic, Chi-squared tests, control vs. FSS-1 week, p = 0.1260, FSS-1 week vs. FSS-2 week, p = 0.0616, n = 10-28, Figure 9i-l).

Consistent with our overall conclusion that irregularly firing ChIs are more sensitive to CRF, we observed differential responses to CRF between the NAc core and shell of control animals (Figure S3) and within the NAc core data set when comparing control animals to FSS-1 week and FSS 2-week animals, particularly at the 10 nM concentration (two-Way RM ANOVA, time x treatment interaction, F_29.319_ = 2.816, p < 0.0001, n = 4-7). We did not see any differences in the CRF response for 3 or 100 nM concentrations. We analyzed the average CRF effect in the last 3 minutes of the experiment for each condition; while there was a trend for a main effect of stress treatment, this was not significant, likely due to the lack of effect at 100 nM and variability in response at 3 nM (two-way ANOVA, main effect of stress treatment, F_2,35_ = 2.912, p = 0.0676, n = 3-8). In contrast to the NAc core, there were no appreciable differences in the CRF potentiation of ChI firing for any concentration we tested when either looking at the time course or average response (same analyses as above, p-values > 0.05, n = 2-10).

## Discussion

In this study, we characterize the spontaneous firing properties of ChIs within the NAc shell of male and female mice. Previous foundational work described three modes of spontaneous firing patterns in dorsal striatal ChIs in 3-4-week-old rats: regular, rhythmic bursting, and irregular (Bennett and Wilson, 1999). Here, we have extended this work to demonstrate that these patterns are conserved across species, sex, striatal regions and well into adulthood using an ex vivo slice preparation. However, we identify a novel fourth category of ChIs that is quantitatively and qualitatively distinct from the other three. This fourth group, which we have named mixed mode, has canonical tonic firing properties like regular or irregular ChIs with intermittent bursting-like rhythmic activity overlaid onto the tonic firing patterns. We further characterized the effect of CRF on the four types of ChI firing patterns and discovered differential sensitivity to CRF, with irregular ChIs potentiating their firing at a the lowest and highest concentration of CRF compared to the other groups along with a marked reduction in the IEI CV. A key insight of this study is that acute CRF application shifts the ChI firing pattern distribution toward more regular firing. We show for the first time that sex and estrous cycle impact the ChI firing rate and proportion of ChI firing patterns, and that there are sex-cycle dependent differences in CRF sensitivity. Finally, we show that repeated stressor exposure increases the heterogeneity of firing pattern proportions in the NAc core and heightens CRF sensitivity while having no impact on the NAc shell. This work is among the first characterizing the firing modes of ChIs in the NAc shell, which may greatly add to our understanding of how these interneurons function on a fundamental level.

Based on previous literature and our own work, these spontaneous firing patterns are intrinsic to the cell since they cannot be altered by blockade of glutamatergic or GABAergic synaptic transmission (Bennett and Wilson, 1999). The difference in firing properties is likely due to a difference in ion channel expression and conductance that is highly dependent on different potassium channel conductances. We and others have shown that attenuation or blockade of HCN channels, sK channels or different Kv channels can cause profound changes in the firing rate, IEI CV and firing pattern distribution and heterogeneity (Bennett, Callaway and Wilson, 2000; Goldberg and Wilson, 2005; Song et al., 1998; Wilson, 2005). For example, attenuation or blockade of sK channels produces a large shift in the CV and firing pattern distribution toward entirely rhythmic bursting cells. These manipulations provide some insight into how endogenous firing modes may be generated and altered. Importantly, Bennett and Wilson (1999) demonstrated that these same spontaneous firing patterns are apparent in the intact animal using extracellular recordings, suggesting that differences in firing motifs may impact behavioral function (Bennett and Wilson, 1999). Without having access to the original data, it is challenging to know for certain why Bennett and Wilson (1999) did not detect the mixed mode ChI type. Based on their reports and our previous work, ChIs recorded in the dorsal striatum tend to have lower firing rates than ChIs in the NAc core or shell (Bennett and Wilson, 1999; Lemos, Shin and Alvarez, 2019), which might make it more difficult to distinguish irregular and mixed mode ChIs. It is also possible that NAc shell ChIs have a different composition of ion channels or sK subtypes within the sK channel family that lead to the emergence of this fourth ChI type. A potential future direction could entail utilizing a method such as PatchSeq to connect ion channel composition and expression for each cell to their firing pattern.

### Neuromodulation and plasticity of ion channel function

Here we report two key findings. First, irregular firing ChIs, characterized by their low firing rate, high CV and lack of bursting, are most responsive to CRF. Second, high concentrations of CRF are able to shift the firing pattern distribution toward more regular firing. CRF exerts its actions on cell excitability via sK channel enhancement in ChIs (Lemos, Shin and Alvarez, 2019); in ChIs it does so by reducing spike accommodation. High sK channel activation gives rise to the medium-duration afterhyperpolarization potential (mAHP), which allows for regular single spiking activity, while low sK channel activity is necessary for bursting (Goldberg and Wilson, 2005). Based on both Lemos et al., (2019) and Goldberg and Wilson (2015), CRF increases sK channel activity in rhythmic bursting and irregular cell types to promote pacemaker activity and prevent bursting activity. High baseline sK activation in regular spiking neurons could occlude CRF effects, particularly at lower concentrations, because regular spiking neurons are already highly active. Though we did not find any studies assessing changes in sK activation across the estrous cycle, changes in the function of other potassium channels across estrous cycle have been reported. In both GnRH neurons of the medial preoptic area and CRF neurons within the paraventricular nucleus of the hypothalamus, I_A_ channel conductance is elevated during estrus and diestrus compared to proestrus (Arroyo et al., 2011; Power and Iremonger, 2021). Regulation of potassium channel conductances may be a conserved mechanism in which the ovarian hormonal cycle can induce transient changes in the excitability of several cell types across the brain.

### Plasticity in ChI firing pattern and CRF modulation driven by sex and estrous cycle

Here we found that NAc shell ChI firing rates and firing patterns differ between male and female mice. While this difference in ChIs is novel, sex differences in the firing rate of other tonically active neurons such as dopamine neurons have been observed (Calipari et al., 2017). This finding indicates to us that there is state-dependent plasticity in spontaneous firing patterns in NAc shell ChIs. Differences in firing properties across the estrous cycle have also been observed in a number of cell types within the mesocorticolimbic pathway. Estrous-cycle driven changes have also been observed in dopamine neuron firing properties (Calipari et al., 2017; Shanley et al., 2023).

Furthermore, in the cortex, there is a pronounced difference in the firing rate and firing pattern of fast-spiking interneurons (FSIs) recorded in awake behaving non-estrus and estrus females. (Clemens et al., 2019). We posit that different modes of firing across estrous alters the network within the brain region to be more or less responsive to environmental stimuli. For example, in the cortex, these FSIs are only responsive to social touch in non-estrus females, not estrus females(Clemens et al., 2019).

Differences in CRF responsivity across estrous also has been observed; however, these have often been in the opposite direct to our own observations (Nappi and Rivest, 1995; Wiersielis et al., 2016). For example, intracerebroventricular administration of CRF increases grooming behavior more in proestrus females compared to diestrus females or males (Wiersielis et al., 2016). Here, we found that a low concentration of CRF (3 nM) can potentiate ChI firing in slices taken from diestrus females, but not estrus females. Novelty has been shown to trigger CRF release in the NAc (Lemos et al., 2012). Differences in CRF sensitivity across the estrous cycle, particularly low concentrations CRF that may encode novelty or salience, but not necessarily stressful/alarming stimuli, may reflect the need to allocate motivational resources differentially throughout the estrous cycle. For example, in diestrus, when reproductive drive is low, females may engage with other environmental stimuli more than during estrus, when reproductive drive is high. While this is speculative, it has been shown that CRF microinfusion into the NAc shell (but not core) increases locomotor behavior (Holahan, Kalin and Kelley, 1997) and that females in diestrus show enhanced locomotor responses to a novel environment compared to females in estrus (Davis et al., 2008; Sell et al., 2005).

### Stress-induced changes in ChI firing properties in the NAc core, but not NAc shell

In this study we found a profound difference in ChI firing properties and firing patterns between the NAc core and shell of male mice. Under control conditions, ChIs in the NAc core displayed a more homogenous firing pattern than the NAc shell since regular firing patterns predominated. Repeated stressor exposure transiently shifted NAc core ChI firing patterns (reflected by a change in CV) to resemble the NAc shell (i.e., more heterogenous, with multiple patterns that dominate including irregular, mixed mode and rhythmic neurons). Over time the firing pattern proportions and CV reverted to the control condition. In contrast, the NAc shell, with cells displaying more heterogenous firing patterns, was not affected by repeated stressor exposure. Again, consistent with our conclusion that irregular ChI neurons are more responsive to CRF, conditions in which more irregular neurons were part of the ChI firing pattern distribution were associated with greater responsivity to CRF. Specifically, in control mice, NAc shell ChIs had higher sensitivity to CRF (10 nM) than NAc core; comparing control NAc core ChIs to FSS-1 week NAc core ChIs revealed a similar enhancement of CRF potentiation of ChI firing. This effect is possibly a combination of a change in the proportions of different firing patterns and modest upregulation of CRF-R1 in ChIs. We propose that repeated stressor exposure transiently creates a window in which NAc core ChIs may be more responsive to salient stimuli in the environment that are encoded by CRF. This may manifest as behaviors such as reduced locomotor habituation to novel environment as seen in Figure 8.

### Limitations

A caveat to our clustering method is the limited information that can be extracted from cell-attached recordings compared to whole-cell current clamp recordings. Future studies should combine the two methods to refine our understanding of ChI firing heterogeneity. Rhythmic bursting cells were sparse in number and were also more challenging to hold at a gigaohm seal through the experiment. This made it challenging to accumulate robust sample sizes for all concentrations of CRF. Thus, we had a significant amount of variability at lower CRF concentrations. Likewise, it was very difficult to capture females in the proestrus and metestrus phases of the estrous cycle since those stages are much shorter than diestrus and estrus. As such, our sample sizes are smaller for those cycle stages. However, considering the difference in the proportions of ChI firing patterns across all four stages, we felt it was important not to pool data into high and low estradiol groups, for example. We can only speculate about what these cellular observations mean for the behaving animal. However, we hope that these ex vivo findings will trigger a new set of testable hypotheses for in vivo studies.

## Acknowledgements

This study was funded by K99/R00 Pathway to Independence award (MH109627) to JCL and NIMH BRAINS R01 (MH122749). We thank Drs. David Lovinger, Veronica Alvarez, and Erin Calipari for their useful input. We also thank Dr. Alvarez for use of her equipment for some of the experiments. Dr. Kavya Devarakonda contributed key edits to the manuscript. The authors declare no competing financial interests.

## Author Contributions

AEI and JCL conceived and designed the research. AEI, YAC, JAS and JCL performed experiments, analyzed data, and interpreted the results of experiments. AEI and JCL prepared the figures, drafted the manuscript, and edited and revised the manuscript. AEI and JCL approved the final version of the manuscript.

## References

1. Al-Hasani, R., Gowrishankar, R., Schmitz, G.P., Pedersen, C.E., Marcus, D.J., Shirley, S.E., Hobbs, T.E., Elerding, A.J., Renaud, S.J., Jing, M., Li, Y., Alvarez, V.A., Lemos, J.C., Bruchas, M.R., 2021. Ventral tegmental area GABAergic inhibition of cholinergic interneurons in the ventral nucleus accumbens shell promotes reward reinforcement. Nat Neurosci 24, 1414–1428.

2. Alonso-Caraballo, Y., Ferrario, C.R., 2019. Effects of the estrous cycle and ovarian hormones on cue-triggered motivation and intrinsic excitability of medium spiny neurons in the Nucleus Accumbens core of female rats. Hormones and behavior 116, 104583.

3. Arroyo, A., Kim, B.S., Biehl, A., Yeh, J., Bett, G.C., 2011. Expression of kv4.3 voltage-gated potassium channels in rat gonadotrophin-releasing hormone (GnRH) neurons during the estrous cycle. Reprod Sci 18, 136–144.

4. Bale, T.L., Vale, W.W., 2004. CRF and CRF receptors: role in stress responsivity and other behaviors. Annual review of pharmacology and toxicology 44, 525–557.

5. Bennett, B.D., Callaway, J.C., Wilson, C.J., 2000. Intrinsic membrane properties underlying spontaneous tonic firing in neostriatal cholinergic interneurons. J Neurosci 20, 8493–8503.

6. Bennett, B.D., Wilson, C.J., 1999. Spontaneous activity of neostriatal cholinergic interneurons in vitro. J Neurosci 19, 5586–5596.

7. Bolam, J.P., Wainer, B.H., Smith, A.D., 1984. Characterization of cholinergic neurons in the rat neostriatum. A combination of choline acetyltransferase immunocytochemistry, Golgi-impregnation and electron microscopy. Neuroscience 12, 711–718.

8. Bonsi, P., Cuomo, D., Martella, G., Madeo, G., Schirinzi, T., Puglisi, F., Ponterio, G., Pisani, A., 2011. Centrality of striatal cholinergic transmission in Basal Ganglia function. Front Neuroanat 5, 6.

9. Calipari, E.S., Juarez, B., Morel, C., Walker, D.M., Cahill, M.E., Ribeiro, E., Roman-Ortiz, C., Ramakrishnan, C., Deisseroth, K., Han, M.H., Nestler, E.J., 2017. Dopaminergic dynamics underlying sex-specific cocaine reward. Nat Commun 8, 13877.

10. Chen, Y.W., Rada, P.V., Butzler, B.P., Leibowitz, S.F., Hoebel, B.G., 2012. Corticotropin-releasing factor in the nucleus accumbens shell induces swim depression, anxiety, and anhedonia along with changes in local dopamine/acetylcholine balance. Neuroscience 206, 155–166.

11. Cheng, J., Umschweif, G., Leung, J., Sagi, Y., Greengard, P., 2019. HCN2 Channels in Cholinergic Interneurons of Nucleus Accumbens Shell Regulate Depressive Behaviors. Neuron 101, 662–672 e665.

12. Clemens, A.M., Lenschow, C., Beed, P., Li, L., Sammons, R., Naumann, R.K., Wang, H., Schmitz, D., Brecht, M., 2019. Estrus-Cycle Regulation of Cortical Inhibition. Curr Biol 29, 605–615 e606.

13. Cora, M.C., Kooistra, L., Travlos, G., 2015. Vaginal Cytology of the Laboratory Rat and Mouse: Review and Criteria for the Staging of the Estrous Cycle Using Stained Vaginal Smears. Toxicol Pathol 43, 776–793.

14. Dabrowska, J., Hazra, R., Guo, J.D., Dewitt, S., Rainnie, D.G., 2013. Central CRF neurons are not created equal: phenotypic differences in CRF-containing neurons of the rat paraventricular hypothalamus and the bed nucleus of the stria terminalis. Front Neurosci 7, 156.

15. Davis, B.A., Clinton, S.M., Akil, H., Becker, J.B., 2008. The effects of novelty-seeking phenotypes and sex differences on acquisition of cocaine self-administration in selectively bred High-Responder and Low-Responder rats. Pharmacol Biochem Behav 90, 331–338.

16. Ding, J.B., Guzman, J.N., Peterson, J.D., Goldberg, J.A., Surmeier, D.J., 2010. Thalamic gating of corticostriatal signaling by cholinergic interneurons. Neuron 67, 294–307.

17. Goldberg, J.A., Wilson, C.J., 2005. Control of spontaneous firing patterns by the selective coupling of calcium currents to calcium-activated potassium currents in striatal cholinergic interneurons. J Neurosci 25, 10230–10238.

18. Gonzales, K.K., Smith, Y., 2015. Cholinergic interneurons in the dorsal and ventral striatum: anatomical and functional considerations in normal and diseased conditions. Ann N Y Acad Sci 1349, 1–45.

19. Hanada, Y., Kawahara, Y., Ohnishi, Y.N., Shuto, T., Kuroiwa, M., Sotogaku, N., Greengard, P., Sagi, Y., Nishi, A., 2018. p11 in Cholinergic Interneurons of the Nucleus Accumbens Is Essential for Dopamine Responses to Rewarding Stimuli. eNeuro 5.

20. Henckens, M.J., Deussing, J.M., Chen, A., 2016. Region-specific roles of the corticotropin-releasing factor-urocortin system in stress. Nat Rev Neurosci 17, 636–651.

21. Holahan, M.R., Kalin, N.H., Kelley, A.E., 1997. Microinfusion of corticotropin-releasing factor into the nucleus accumbens shell results in increased behavioral arousal and oral motor activity. Psychopharmacology (Berl) 130, 189–196.

22. Itoga, C.A., Chen, Y., Fateri, C., Echeverry, P.A., Lai, J.M., Delgado, J., Badhon, S., Short, A., Baram, T.Z., Xu, X., 2019. New viral-genetic mapping uncovers an enrichment of corticotropin-releasing hormone-expressing neuronal inputs to the nucleus accumbens from stress-related brain regions. J Comp Neurol 527, 2474–2487.

23. Krok, A.C., Maltese, M., Mistry, P., Miao, X., Li, Y., Tritsch, N.X., 2023. Intrinsic dopamine and acetylcholine dynamics in the striatum of mice. Nature.

24. Land, B.B., Bruchas, M.R., Lemos, J.C., Xu, M., Melief, E.J., Chavkin, C., 2008. The dysphoric component of stress is encoded by activation of the dynorphin kappa-opioid system. J Neurosci 28, 407–414.

25. Lemos, J.C., Shin, J.H., Alvarez, V.A., 2019. Striatal cholinergic interneurons are a novel target of corticotropin releasing factor. J Neurosci.

26. Lemos, J.C., Wanat, M.J., Smith, J.S., Reyes, B.A., Hollon, N.G., Van Bockstaele, E.J., Chavkin, C., Phillips, P.E., 2012. Severe stress switches CRF action in the nucleus accumbens from appetitive to aversive. Nature 490, 402–406.

27. Lim, M.M., Liu, Y., Ryabinin, A.E., Bai, Y., Wang, Z., Young, L.J., 2007. CRF receptors in the nucleus accumbens modulate partner preference in prairie voles. Hormones and behavior 51, 508–515.

28. Mendonca, P.R., Vargas-Caballero, M., Erdelyi, F., Szabo, G., Paulsen, O., Robinson, H.P., 2016. Stochastic and deterministic dynamics of intrinsically irregular firing in cortical inhibitory interneurons. Elife 5.

29. Mendonca, P.R.F., Kyle, V., Yeo, S.H., Colledge, W.H., Robinson, H.P.C., 2018. Kv4.2 channel activity controls intrinsic firing dynamics of arcuate kisspeptin neurons. J Physiol 596, 885–899.

30. Merali, Z., McIntosh, J., Anisman, H., 2004. Anticipatory cues differentially provoke in vivo peptidergic and monoaminergic release at the medial prefrontal cortex. Neuropsychopharmacology 29, 1409–1418.

31. Mohebi, A., Collins, V.L., Berke, J.D., 2023. Accumbens cholinergic interneurons dynamically promote dopamine release and enable motivation. Elife 12.

32. Nappi, R.E., Rivest, S., 1995. Ovulatory cycle influences the stimulatory effect of stress on the expression of corticotropin-releasing factor receptor messenger ribonucleic acid in the paraventricular nucleus of the female rat hypothalamus. Endocrinology 136, 4073–4083.

33. Nosaka, D., Wickens, J.R., 2022. Striatal Cholinergic Signaling in Time and Space. Molecules 27.

34. Oswald, M.J., Oorschot, D.E., Schulz, J.M., Lipski, J., Reynolds, J.N., 2009. IH current generates the afterhyperpolarisation following activation of subthreshold cortical synaptic inputs to striatal cholinergic interneurons. J Physiol 587, 5879–5897.

35. Pecina, S., Schulkin, J., Berridge, K.C., 2006. Nucleus accumbens corticotropin-releasing factor increases cue-triggered motivation for sucrose reward: paradoxical positive incentive effects in stress? BMC biology 4, 8.

36. Power, E.M., Iremonger, K.J., 2021. Plasticity of intrinsic excitability across the estrous cycle in hypothalamic CRH neurons. Sci Rep 11, 16700.

37. Razidlo, J.A., Fausner, S.M.L., Ingebretson, A.E., Wang, L.C., Petersen, C.M., Mirza, S., Swank, I.N., Alvarez, V.A., Lemos, J.C., 2022. Chronic loss of muscarinic M5 receptor function manifests disparate impairments in exploratory behavior in male and female mice despite common dopamine regulation. J Neurosci 42, 6917–6930.

38. Sell, S.L., Dillon, A.M., Cunningham, K.A., Thomas, M.L., 2005. Estrous cycle influence on individual differences in the response to novelty and cocaine in female rats. Behav Brain Res 161, 69–74.

39. Shanley, M.R., Miura, Y., Guevara, C.A., Onoichenco, A., Kore, R., Ustundag, E., Darwish, R., Renzoni, L., Urbaez, A., Blicker, E., Seidenberg, A., Milner, T.A., Friedman, A.K., 2023. Estrous Cycle Mediates Midbrain Neuron Excitability Altering Social Behavior upon Stress. J Neurosci 43, 736–748.

40. Song, W.J., Tkatch, T., Baranauskas, G., Ichinohe, N., Kitai, S.T., Surmeier, D.J., 1998. Somatodendritic depolarization-activated potassium currents in rat neostriatal cholinergic interneurons are predominantly of the A type and attributable to coexpression of Kv4.2 and Kv4.1 subunits. J Neurosci 18, 3124–3137.

41. van de Velden, M., Iodice D’Enza, A. and Markos, A., 2018. Distance-based clustering of mixed data. WIREs Computational Statistics.

42. Warner-Schmidt, J.L., Schmidt, E.F., Marshall, J.J., Rubin, A.J., Arango-Lievano, M., Kaplitt, M.G., Ibanez-Tallon, I., Heintz, N., Greengard, P., 2012. Cholinergic interneurons in the nucleus accumbens regulate depression-like behavior. Proc Natl Acad Sci U S A 109, 11360–11365.

43. Wiersielis, K.R., Wicks, B., Simko, H., Cohen, S.R., Khantsis, S., Baksh, N., Waxler, D.E., Bangasser, D.A., 2016. Sex differences in corticotropin releasing factor-evoked behavior and activated networks. Psychoneuroendocrinology 73, 204–216.

44. Wilson, C.J., 2005. The mechanism of intrinsic amplification of hyperpolarizations and spontaneous bursting in striatal cholinergic interneurons. Neuron 45, 575–585.

45. Zhang, Y.F., Cragg, S.J., 2017. Pauses in Striatal Cholinergic Interneurons: What is Revealed by Their Common Themes and Variations? Front Syst Neurosci 11, 80.

46. Zucca, S., Zucca, A., Nakano, T., Aoki, S., Wickens, J., 2018. Pauses in cholinergic interneuron firing exert an inhibitory control on striatal output in vivo. Elife 7.

