## Supplemental Materials for "Corticotropin releasing factor alters the functional diversity of accumbal cholinergic interneurons"

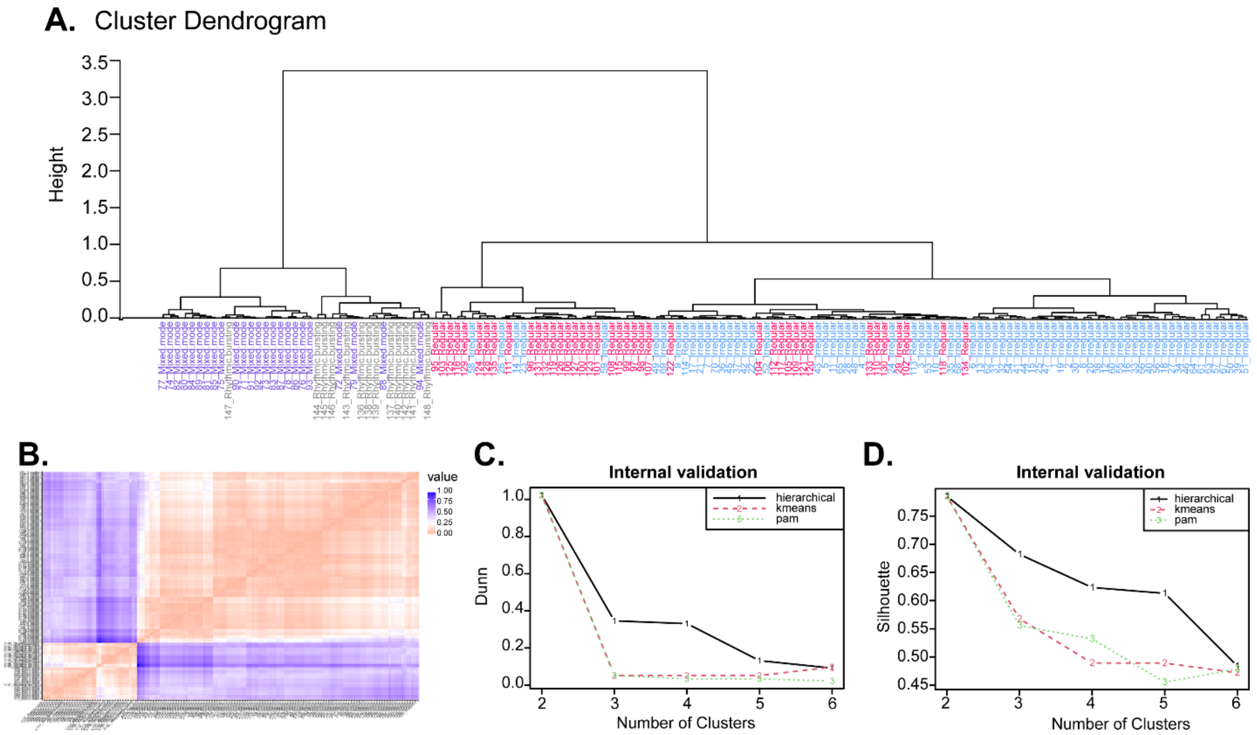


**Figure S1, related to Figure 1. Hierarchical clustering of baseline ChI with internal cluster validation.** A) Cluster dendogram based on hierarchical clustering model of 145 ChIs. The experimenter assigned classification is shown below (regular = magenta, irregular = light blue, rhythmic bursting = grey, mixed mode = purple). B) Dissimilarity matrix, calculated using the *daisy* function in the R cluster package to calculate the Gower distance, a distance metric appropriate for scaling mixed variable types, including continuous and categorical. C) Dunn analysis on three different clustering models, hierarchical (solid black), k-means (red dash), pam (green dash) indicating the optimal number of clusters. D) Silhouette coefficient on three different clustering models, hierarchical (solid black), k-means (red dash), pam (green dash) indicating the optimal number of clusters.

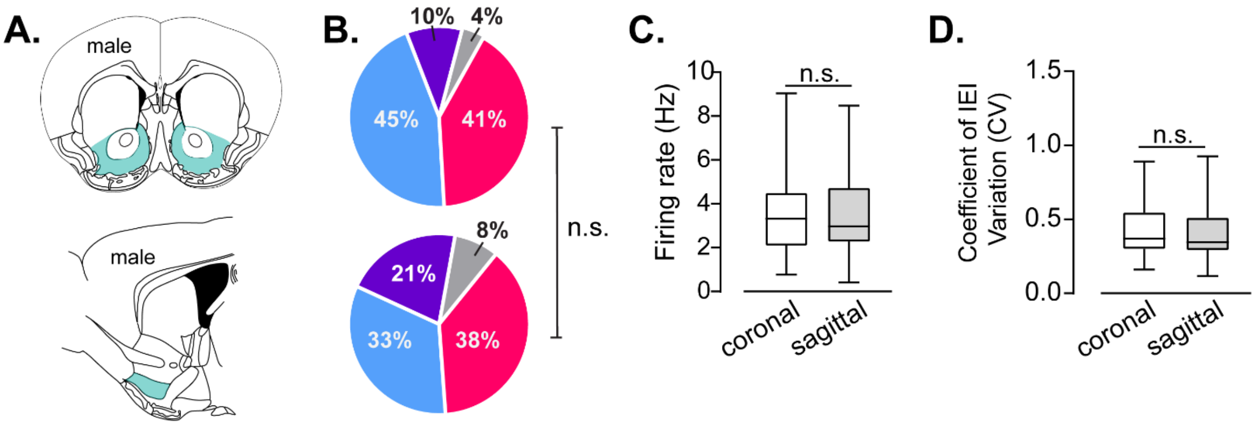


**Figure S2, related to Figure 9. Plane of section does not significantly change ChI firing pattern distribution.** A) Schematic showing a typical location of NAc shell recordings in the coronal (top) or sagittal (bottom) plane. B) ChI firing pattern distribution of ChIs recorded from NAc shell of male mice in the coronal (top) or sagittal (bottom) plane of section. n.s. = Chi-squared test, $p > 0.05$. (regular = magenta, irregular = light blue, rhythmic bursting = grey, mixed mode = purple). C) Mean firing rate of ChIs recorded from NAc shell of male mice in the coronal or sagittal plane of section. Mann-Whitney test, $p = 0.8909$. D) Mean CV of IEI of ChIs recorded from NAc shell of male mice in the coronal or sagittal plane of section. Mann-Whitney test, p = 0. 6868. N = 30 coronal ChIs, 23 sagittal ChIs.

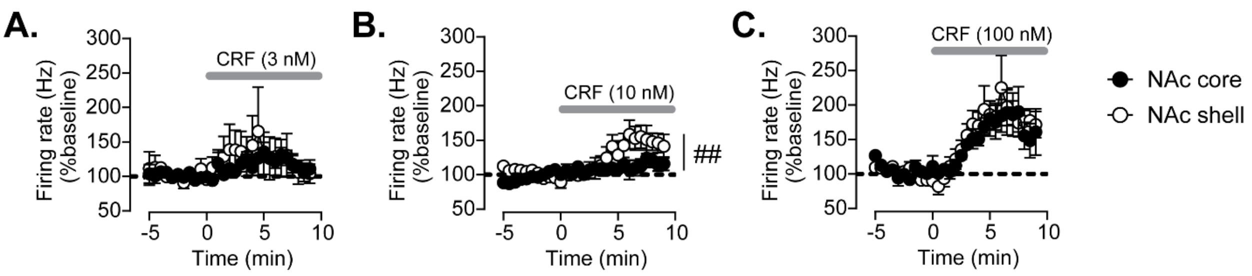


**Figure S3, related to Figure 9. Region-dependent differences in CRF potentiation of ChI firing in control male mice.** A) Timecourse of normalized firing rate (% baseline) prior to and following bath application of 3 nM CRF in ChIs recorded from the NAc core or NAc shell of control mice. B) Timecourse of normalized firing rate (% baseline) prior to and following bath application of 10 nM CRF in ChIs recorded from the NAc core or NAc shell of control mice. ## = Two-Way RM ANOVA, time x stress treatment interaction, $F_{29,348} = 1.965$, $p = 0.0026$. C) Timecourse of normalized firing rate (% baseline) prior to and following bath application of 100 nM CRF in ChIs recorded from the NAc core or NAc shell of control mice.
